# Microglial Stim1/Stim2 deletion reduces calcium signaling but promotes infarct expansion after ischemic stroke

**DOI:** 10.64898/2026.11.12.766275

**Authors:** Khadijeh A. Sharifi, Pedro Norat, Nina Jannatifar, Ziyuan Wang, Carissa Vergeres, Misha Padigala, M. Filip Sluzewski, Lei Liu, Anran Zhao, Matthew R. Hoch, Hao Le, Maurits A. Jansen, Ryan T. Kellogg, Chia-Yi Kuan, G. Wilson Miller, Petr Tvrdik

## Abstract

Microglia respond to ischemic stroke with calcium transients carried by calcium release-activated calcium (CRAC) channels, gated by the endoplasmic reticulum sensors Stim1 and Stim2. Since CRAC signaling drives pro-inflammatory responses in immune cells, blocking it in microglia should protect the ischemic brain. We have examined this hypothesis in 15-month-old mice lacking *Stim1* and *Stim2* in microglia (*Cx3cr1-*CreERT2; *Stim* cKO). In vivo two-photon imaging confirmed that cKO microglia produced one-third smaller calcium transients during cortical spreading depolarizations. We then followed distal middle cerebral artery occlusion (dMCAo) with multiparametric MRI at 9.4 T. Six hours after occlusion, both the control and cKO genotypes demonstrated similar infarct size, edema, and diffusion, but lesion perfusion tended to be lower in cKO mice. Differences in tissue injury emerged later. Between 6 and 48 hours, the infarcts in control males did not grow, whereas infarcts in cKO males expanded 2.7-fold. At 48 hours, cKO males had 2.2-fold larger infarcts, while cKO females did not differ from controls. Moreover, cKO lesions showed a lower apparent diffusion coefficient, indicating more severe tissue injury. Neither hemispheric swelling nor cerebral arterial anatomy accounted for these differences. Contrary to our hypothesis, ablating microglial CRAC channels worsened stroke outcome. Microglial CRAC signaling may therefore protect threatened tissue in the first days after ischemia, possibly by sustaining its perfusion, and this role appears to outweigh its pro-inflammatory effects.

## INTRODUCTION

Stroke is the second leading cause of death and a leading cause of long-term disability worldwide. Ischemic stroke, in which occlusion of a cerebral artery reduces blood flow and causes hypoxic tissue damage, accounts for 80–90% of cases (Barthels & Das, 2020; Henderson et al., 2018). The resulting energy deficit disrupts ionic homeostasis, causing neurons to lose their ability to maintain membrane potential. This ionic dysregulation frequently triggers progressive cortical spreading depolarizations (CSDs)—synchronized waves of neuronal and glial depolarization that propagate across the cortex from the site of injury (Ayata & Lauritzen, 2015; Kearns et al., 2022; Lauritzen et al., 2011; Wainsztein & Rodriguez Lucci, 2018). CSDs exacerbate ischemic damage by increasing metabolic demand in already compromised tissue, further depleting energy reserves and expanding the area of injury (Dreier, 2011; Hartings, 2017).

Microglia originate from primitive yolk sac myeloid progenitors during embryonic development (Ginhoux et al., 2010). However, in adulthood, however, they are maintained within the brain parenchyma independently of postnatal hematopoiesis (Prinz et al., 2021). Under homeostatic conditions, microglia continuously survey their microenvironment with their highly motile processes, making direct contact with neurons, astrocytes, pericytes, and endothelial cells within the neurovascular unit (NVU) (Cserep et al., 2021; Lenart et al., 2024). Notably, microglia are among the earliest responders to ischemic injury and exhibit rapid morphological changes and increased process motility within minutes to hours after the insult (Davalos et al., 2005; Szalay et al., 2016). Upon activation, microglia transition from a ramified, surveillant phenotype to an amoeboid morphology, engage in phagocytosis, and release both pro-inflammatory cytokines (TNF-α, IL-1β, IL-6) and anti-inflammatory mediators (IL-10, TGF-β, Arg-1). The balance of these responses is shaped by signals from the surrounding microenvironment (Qin et al., 2019; Wang et al., 2023).

Microglia play complex, sometimes opposing roles in ischemic stroke. Several studies have demonstrated that pharmacological depletion of microglia using CSF1R inhibitors or genetic ablation strategies leads to larger infarct volumes and worse neurological outcomes, suggesting an important neuroprotective function (Marino Lee et al., 2021; Szalay et al., 2016). The neuroprotective role is supported by evidence showing that microglial P2Y12 receptor signaling is crucial for maintaining cerebrovascular reactivity. Blocking this signaling results in larger lesion volumes, increased neuronal calcium activity, and poorer neurological outcomes (Csaszar et al., 2022; Cserép et al., 2020). Conversely, selective microglial depletion at early stages after photothrombotic stroke has been shown to reduce infarct volume, decrease neurodegeneration, and improve behavioral outcomes (Li et al., 2021). These conflicting findings may reflect differences in stroke models, depletion methods, timing, and the stage-dependent nature of microglial phenotypes (Lenart et al., 2024; Wang et al., 2023).

Previous studies have found that microglia play a role in the initiation and progression of CSDs (Szalay et al., 2016). Consistently, we have previously demonstrated that focal cerebral ischemia caused by middle cerebral artery occlusion (MCAo) in mice results in recurrent CSDs, which in turn lead to frequent calcium transients in microglia (Liu et al., 2021). These damage-induced calcium transients (DICTs) are specific to microglia and contribute to their morphological response to tissue injury, which is mediated by ATP binding to P2Y receptors and requires calcium release from intracellular stores (Hasan et al., 2024). Intracellular calcium (Ca²⁺) signaling is a crucial regulatory pathway in microglia, governing essential cellular functions including migration, phagocytosis and cytokine production (Hasan et al., 2024). Importantly, our study found that CSD-associated microglial calcium activity is due to calcium influx through calcium-release-activated-calcium (CRAC) channels (Liu et al., 2021).

CRAC channel signaling is mediated by the store-operated calcium entry (SOCE) pathway, in which stromal interaction molecules (STIMs), including Stim1 and Stim2, serve as calcium sensors on the endoplasmic reticulum (ER) membrane (Hasan et al., 2024). When intracellular Ca²⁺ stores become depleted, STIM proteins undergo structural alterations and physically interact with Orai channel proteins on the plasma membrane, forming the CRAC channel complex that allows extracellular calcium to enter the cell (Prakriya & Lewis, 2015). This calcium influx activates calcineurin, which dephosphorylates the nuclear factor of activated T cells (NFAT) transcription factors, leading to the transcription of inflammatory mediators, including IL-1β and TNF-α (Mizuma et al., 2019). In the context of stroke, microglial secretion of TNF-α has been shown to lower the threshold for CSD induction, creating a positive feedback loop whereby CSD-triggered microglial calcium signaling promotes further TNF-α release that perpetuates CSD propagation (Grinberg et al., 2013; Grinberg et al., 2012; Jander et al., 2001). This mechanistic link between CRAC channel-mediated calcium signaling and neuroinflammation led us to predict that deletion of CRAC channel components would improve stroke outcome by reducing acute post-stroke neuroinflammation.

In this study, we tested this hypothesis by generating mice with microglia-specific conditional deletion of both *Stim1* and *Stim2* genes, thereby ablating functional CRAC channels in microglia. Unexpectedly, although deletion of both Stim sensors effectively reduced microglial calcium signaling, it resulted in more severe stroke outcomes following MCAo. These findings suggest that CRAC channel-mediated calcium signaling in microglia serves additional beneficial functions during acute ischemic injury that extend beyond their well-characterized pro-inflammatory role, and that the neuroprotective contributions of microglial calcium signaling may outweigh its detrimental inflammatory effects in the hyperacute phase of stroke.

## METHODS

### Animal models and surgical procedures for ischemic stroke

The generation of the Cre-dependent GCaMP5G- and tdTomato-expressing reporter (PC::G5-tdT) has been described (Gee et al., 2014). *Cx3cr1*-*CreERT2* mice (Yona et al., 2013) were purchased from The Jackson Laboratory. *Stim1* and *Stim2* floxed mice were provided by Dr. Stefan Feske (Oh-hora & Rao, 2008). Experimental mutants were homozygous for both floxed *Stim* alleles and carried *Cx3cr1-CreERT2* and PC::G5-tdT; littermate controls were heterozygous for the floxed *Stim* alleles and carried the same Cre and reporter. Both sexes were used. At 10 months of age, all mice received tamoxifen for five consecutive days to induce recombination in CX3CR1-expressing cells. Experiments were performed at 15 months of age. Focal cerebral ischemia was induced by permanent occlusion of the middle cerebral artery.

Proximal occlusion used silicon-coated filaments (Tvrdik et al., 2019); distal occlusion (dMCAo) was performed as described (Norat et al., 2023) using Spetzler Malis Forceps, 7”, 1.0 mm (Stryker SKR6760180010) (Figure S3). The 6-hour and 48-hour dMCAo studies were independent cohorts, and no animal was imaged at both time points. Animals were randomized before surgery or imaging, and the surgeon was blinded to genotype. The 6-hour cohort was restricted to males, whereas the 48-hour cohort included males and females. All procedures were approved by the University of Virginia Institutional Animal Care and Use Committee.

### Microglia isolation, RNA extraction, cDNA synthesis, and Real-Time PCR

Adult mouse brains were dissected and enzymatically dissociated using the Adult Brain Dissociation Kit (Miltenyi Biotec, #130-107-677) according to the manufacturer’s protocol. Cell suspensions were centrifuged to remove debris, and microglia were selected using CD11b MicroBeads (Miltenyi Biotec, #130-049-601). Approximately 200,000–300,000 cells were recovered per brain. Total RNA was extracted using the Quick-RNA Microprep Kit (Zymo Research, #R1050), and cDNA was synthesized from 10 ng of total RNA using the iScript cDNA Synthesis Kit (Bio-Rad, #1708890). We verified the purity of the microglial isolate by PCR with primers for the microglial marker P2ry12, along with the markers for neurons (NeuN, Rbfox3), astrocytes (Gfap), and oligodendrocytes (Olig1). PCR products were resolved on a 2% agarose gel and visualized with the Azure Imaging System. Quantitative PCR was performed on a Bio-Rad CFX96 Real-Time PCR Detection System using iQ SYBR Green Supermix (Bio-Rad, #1708880). Relative gene expression was calculated using the 2^–ΔΔCq method, with *Actb* (β-actin) as the reference gene. The primer sequences for *Stim1*, *Stim2*, *P2ry12* (Fu et al., 2025), *NeuN* (*Rbfox3*), *Gfap*, and *Olig1* (Chen et al., 2022), as well as *Actb* (Pratama et al., 2015) are listed in Table 1.

**Table 1.**
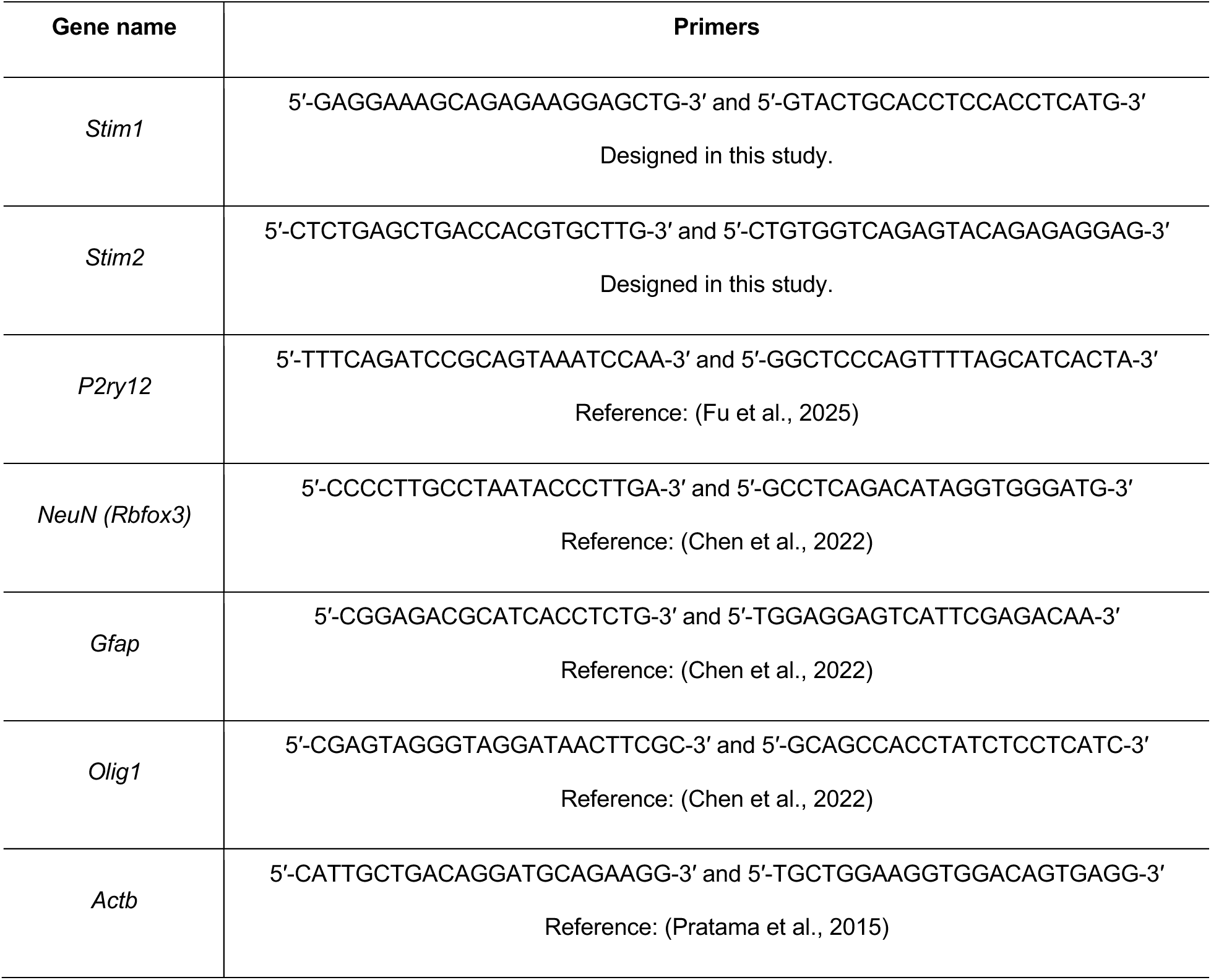
Primer sequences used for RT-PCR.

### Cranial window surgery and two-photon (2p) imaging

In deeply anesthetized animals, the skin and periosteum were removed from the skull and a 3-4 mm diameter circle was drilled in the right parietal bone. The skull flap was gently removed with fine forceps, and a circular glass coverslip was placed over the exposed dura and sealed around the edges with superglue. A metal pin headpost was attached to the parietal bone and the exposed bone was covered with dental cement (Tvrdik et al., 2019). Lightly anesthetized animals were then mounted on the microscope stage, and cortical microglia were imaged with an Olympus FVMPE-RS at 2.5 Hz, 512 x 512 pixel resolution. The pulsed infrared lasers were tuned to 920 nm, and the emitted photons were captured by green- and red-channel detectors.

### Induction of CSD

Chemical induction of CSD was performed through burr holes drilled over the visual cortex, posterior to the cranial window. After mounting the mouse on the imaging platform, a fine-tip glass micropipette filled with 1M KCl was loaded in the programmable Nanoject injector and lowered into the burr hole using a precise micromanipulator. One hundred nL of 1 M KCl was expelled into the cortex 1 min after the initiation of high-speed imaging, and the resulting CSDs were recorded with 1.05 NA, 25X lens, and 2x zoom.

### 2p image processing and analysis

T-series were processed in Imaris 9.5 (Bitplane). Each recording was first inspected, and 3-6 tdTomato-labeled microglial cells with cell bodies in the focal plane were selected for analysis. A dynamic region of interest (ROI) was defined in the red (tdTomato) channel and tracked across the recording, so that the ROI followed the changing shape of each soma independently of calcium-dependent fluorescence. Mean green (GCaMP5G) intensity within each tracked ROI was exported for every time point. Signal processing was performed with custom Python scripts using filters from the SciPy package. Traces were denoised with a mean filter followed by a Wiener filter (kernel 5 timepoints, ≈ 2 s), with noise modeled as the standard deviation of the soma’s mean fluorescence at each timepoint. Because background fluorescence drifted before and after passage of the CSD, baseline was treated as a function of time rather than as a pre-stimulus average: F₀(t) was estimated by applying a median filter (101 timepoints, ≈ 40.4s), which removes the CSD response, followed by a Wiener filter (71 timepoints, ≈ 28.4 s) to suppress residual high-frequency noise. The final trace was calculated as ΔF/F₀ = [F(t) − F₀(t)] / F₀(t). Responses were measured over a window from 60 to 200 s after the start of imaging: peak amplitude was the maximum value within the window; duration was the interval between the last time point below zero preceding the peak and the first time point below zero following it; and area under the curve was integrated over that same interval. The algorithm was written by M. F. Sluzewski; the Jupyter notebook is available on request. Representative traces at each processing stage are shown in Figure S1C–E.

### MRI sequences

All in vivo MRI was performed on a 9.4 T Bruker BioSpec small-animal scanner (ParaVision 360.3.4) with a volume transmit coil and a mouse-brain array receive coil. Mice were anesthetized with 1–2% isoflurane during scanning, and body temperature was maintained at 35–37 °C with a heated cradle. At 6 h and 48 h after occlusion, T2-weighted, diffusion-weighted (DWI) and arterial spin labeling (ASL) sequences were acquired within a single 30–45 min session to assess edema, diffusion restriction and perfusion, respectively; time-of-flight MR angiography was added at 48 h.

T2-weighted images were acquired with a 2D rapid acquisition with relaxation enhancement (TurboRARE) spin-echo sequence (effective echo time 34 ms, turbo factor 8, refocusing flip angle 126–180°, 4 signal averages, matrix 220 × 220, field of view 20 × 20 mm, in-plane pixel 91 × 91 µm, slice thickness 0.5 mm). Coronal stacks had 30–33 contiguous slices (repetition time 2500–2746 ms; 4.5–4.9 min). Axial stacks had 13–18 slices (repetition time 1082–2500 ms; 1.9–4.5 min); they were acquired in both cohorts and segmented at 48 h.

Time-of-flight MR angiography images were acquired with a 3D fast low-angle shot (FLASH) spoiled gradient-echo sequence with off-resonance magnetization transfer preparation (repetition time of 30 ms, echo time of 3.14 ms, excitation flip angle of 30°, readout bandwidth of 306 Hz/pixel, RF- and gradient-spoiled, 75% partial Fourier sampling in the phase-encoding direction, acquisition matrix 140x120x114 reconstructed to 170x160x104, field of view of 16x16x16 mm, reconstructed voxel size of 100x100x154 μm, 2 signal averages, and 13.7 min acquisition time). A spatially selective saturation slab (median thickness 9.5 mm, range 6.1–13.5 mm) was positioned to suppress venous inflow signal.

Diffusion-weighted images were acquired with a segmented spin-echo echo-planar imaging (EPI) sequence (repetition time 2000 ms, effective echo time 16.9 ms, 4 segments with an echo train of 15 and 62.5% partial-Fourier sampling of 96 phase-encoding lines, readout bandwidth 3472 Hz/pixel, 4 averages, 2.1 min). The matrix was 96 × 96 with a 15 × 15 mm field of view (156 µm pixels) at 48 h and in the first 6-h session, and a 20 × 20 mm field of view (208 µm pixels) in the second 6-h session, in 30–33 contiguous coronal slices of 0.5 mm. Each acquisition comprised a reference image (b = 22–24 s/mm^2^, imaging gradients only) and diffusion weighting along three orthogonal directions (b = 690–706 s/mm^2^). The b-value of each image was taken as the trace of the b-matrix recorded by the scanner. Phase encoding was dorso-ventral.

Perfusion was measured with flow-sensitive alternating inversion recovery (FAIR) ASL and a segmented EPI readout (repetition time 10 s, echo time 11 ms, 2 segments with an echo train of 31 and 77.5% partial-Fourier sampling of 80 phase-encoding lines, readout bandwidth 4261 Hz/pixel, 1 average) in a single 1-mm coronal slice through the infarct (matrix 80 × 80, field of view 18 × 18 mm, 225 × 225 µm pixels). Slice-selective and non-selective inversion images were acquired at 13 inversion times (TI = 12, 100, 200, 300, 400, 500, 600, 800, 1200, 1400, 1600, 2000 and 2300 ms; 9.4 min). Scans acquired at the 6-h timepoint contained three additional TIs (4000, 6000 and 8000 ms; 12.6 min). All acquired TIs were used: 16 at 6 h and 13 at 48 h. ASL uses arterial blood water as an endogenous tracer: the inversion labels inflowing blood, and the difference between the two inversion conditions reflects tissue perfusion (Kim, 1995).

### MRI data analysis

Infarct (T2-hyperintense tissue) and whole brain were traced manually on every slice of the T2 stacks in ITK-SNAP (Yushkevich et al., 2006). At 6 h, two investigators traced the coronal stack. At 48 h, two investigators traced both the coronal and the axial stacks. Infarct volume is the mean of the two (6 h) or four (48 h) tracings, and relative infarct volume is the infarct as a percentage of the traced brain volume. Lesions for the primary analyses were not traced on the diffusion or perfusion images; masks that the second investigator drew on the 6-h perfusion maps served only as a sensitivity analysis (Table SM4). All lesion regions for ADC, CBF and T2 signal were derived from the coronal T2 tracings of the investigator who segmented both cohorts; the second investigator’s tracings were used for sensitivity analyses. All further steps used custom Python code, available from the corresponding authors on request; full details are given in the Supplementary Methods.

For DWI, the reference image of each scan was first registered to the T2 stack on which the masks were drawn. The registration combined a 3-D translation, for head motion between series, with a smooth displacement along the phase-encoding axis, for EPI distortion (Jezzard & Balaban, 1995), and maximized normalized cross-correlation. Each DWI voxel (4 × 4 × 3 points) or ASL voxel (5 × 5 × 5 points) was supersampled to obtain the fractions of brain and lesion it contained. Voxels with at least 50% brain formed the brain region, and brain voxels with at least 50% lesion formed the lesion region. In every DWI slice, and in the ASL slab, the mid-sagittal line was fitted by reflection symmetry of the brain outline and the T2 intensity, with the lesion left out of the intensity term. The contralateral region was the lesion reflected across this line, restricted to contralateral brain and stripped of any voxel that an investigator had traced as lesion (Figure S4D).

ADC was computed per voxel as the trace estimate, ADC = Σ ln(S_0_/S_i_) / Σ (b_i_ − b_0_), summed over the diffusion directions (Basser et al., 1994). Segmented EPI without gating is sensitive to motion, which can cancel the signal of a whole slice in one acquisition (Anderson & Gore, 1994). Such dropouts were detected in normal-appearing tissue as slices whose median diffusion-weighted/reference signal ratio, or reference signal, deviated by more than 0.10 (natural logarithm) from its five-slice running median. A slice was used when its reference image and at least two directions were intact, and ADC was then computed from the intact directions only. Lesion and contralateral regions were always evaluated on the same slices and directions. Voxels with ADC ≤ 0 or > 1.5 × 10^−3^ mm^2^/s (cerebrospinal fluid) were excluded. Lesion and contralateral ADC are voxel means pooled over all usable slices that contained both regions, and relative ADC (% ADC) is their ratio × 100. Each animal required at least two usable lesion slices covering at least 50% of the lesion; all animals met this criterion.

CBF was quantified pixel by pixel. The magnitude inversion-recovery model S(TI) = |M_0_ [1 − 2 exp(−TI/T_1_)]| was fitted separately to the slice-selective and non-selective series of each pixel, giving T_1,sel_ and T_1,non_. CBF (ml/100 g/min) = 6000 · λ · (T_1,non_/T_1,blood_) · (1/T_1,sel_ − 1/T_1,non_) (^B^elle et al., 199^8^; ^K^ober et al., 200^8^), with T_1_ in seconds, λ = 0.9 ml/g (^H^erscovitch & Raichle, ^1^98^5^) and T_1,blood_ = 2.43 s (^D^obre et al., 200^7^). Lesion and contralateral CBF are the means of pixel CBF within each region. We retained negative pixel values, which noise produces where flow is low, so that the mean was not biased upwards. We fitted before averaging because averaging magnitude signals across pixels distorts the inversion-recovery curve near its null. In simulations of each 6-h scan with known CBF, the pixel mean was nearly unbiased, whereas a fit of the region-averaged signals was not (Supplementary Methods). Two quality rules, defined on the signals alone, were applied to every scan. First, at TIs of 4000 ms and longer, the slice-selective and non-selective signals have almost fully recovered and should nearly coincide. A long TI was therefore dropped from a scan when the two differed by more than 5% of M_0_ in the lesion or contralateral region. This removed one image pair (6000 ms in one 6-h mouse). Second, pixels whose M_0_ was below ten times the background noise were excluded from all regions (five lesion pixels in two 6-h mice, three contralateral pixels in two 48-h mice). Relative CBF (% CBF) is lesion CBF as a percentage of contralateral CBF. The same model produced the CBF maps in Figure 3A. No CSF exclusion was applied to CBF. Scans in which the contralateral hemisphere had a CBF below 20 ml/100 g/min (absent perfusion) were excluded from CBF analysis.

Relative T2-weighted signal (% T2) was the mean T2-weighted signal in the lesion as a percentage of that in the contralateral region, both defined on the native coronal T2 slices with the per-slice midline described above.

Swelling was measured on the coronal T2 tracings. In each slice, the traced brain (ventricles included) was split at a mid-sagittal line fitted as above, but with the lesion and its mirror image hidden from both the outline and the intensity terms, so that the swollen side could not shift the line. Swelling was 100 × (ipsilateral − contralateral)/contralateral volume within the infarcted slab (lesion slices ± 1 mm). Because every infarct was in the right hemisphere, the mean left–right asymmetry of lesion-free slices (at least 2 mm from the lesion) was subtracted as a constant baseline (+0.7% and +0.8% for the two investigators’ 48-h tracings). At 48 h, the two investigators’ values were averaged; at 6 h, brain outlines were available from one investigator. The swelling-related excess volume, the baseline-corrected difference between ipsilateral and contralateral slab volumes, was subtracted from 48-h infarct volumes for the edema-adjusted comparison (Figure S4C). Axial tracings were not used, because head roll is through-plane in horizontal slices.

When a sequence had been repeated, the scan was chosen before any lesion value was computed. For DWI, the order of preference was coronal orientation, dorso-ventral phase encoding, the most usable lesion slices, the fewest dropouts, and the best registration. For ASL, it was the best inversion-recovery fit quality (10th percentile of the brain R^2^), then the highest M_0_ signal-to-noise ratio.

### Histology

Following the MRI sessions, mice were euthanized with an intraperitoneal injection of Euthasol for histopathological verification of infarct size. Brains were quickly removed and sliced into 1-mm-thick coronal sections using a mouse brain matrix. Slices were incubated in 2,3,5-triphenyltetrazolium chloride (TTC) solution (2% w/v in PBS) at 37 °C for 25 minutes. TTC is a metabolic stain that labels viable tissue red by reacting with NADH-dependent dehydrogenases, whereas infarcted (non-viable) tissue remains pale/white. After staining, slices were fixed in 4% paraformaldehyde for imaging. Each slice was photographed with a digital CCD camera mounted on an Olympus stereomicroscope. The infarct area (white region) was outlined and measured on each section using ImageJ 1.54. TTC-derived infarct volumes (summed across slices) were later compared with MRI-derived volumes for consistency. TTC infarct volumes were compared qualitatively with MRI volumes in the brains with adequate staining.

### Statistical analysis

Data are reported as mean ± SD unless noted otherwise. Tests were two-sided except for the qPCR confirmation of gene deletion. Significance was set at p < 0.05. The test used for each comparison is stated in the corresponding Figure legend. Analyses were performed in GraphPad Prism and in Python 3.13 (NumPy 2.3, SciPy 1.15, statsmodels 0.14, pandas 3.0, nibabel 5.4, pydicom 3.0 and scikit-image 0.26).

*Quantitative RT-PCR* (Figure 1C, D). Relative *Stim1* and *Stim2* expression was calculated by the 2–ΔΔCq method with *Actb* as the reference gene. Groups (n = 3 mice per group) were compared on the ΔCq scale with an unpaired one-tailed Student’s t-test. The one-tailed test was specified in advance because Cre-mediated excision of the floxed exons can only reduce transcript abundance.

**Figure 1.**
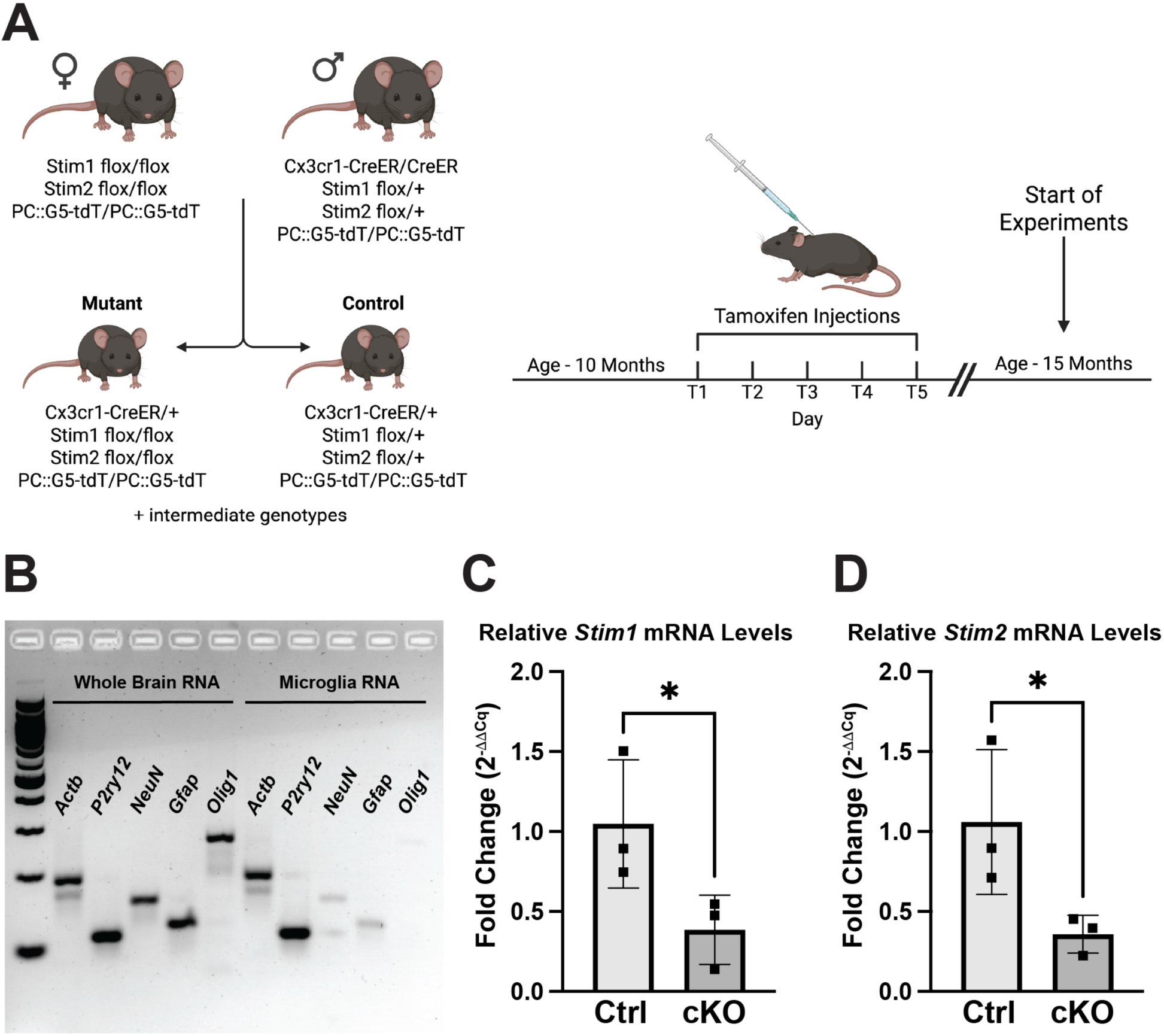
Conditional deletion of Stim1 and Stim2 in microglia and validation of gene knockout. (A) Schematic of the breeding strategy and experimental timeline. Female mice homozygous for floxed *Stim1* and *Stim2* alleles and homozygous for the PC::G5-tdT calcium indicator were crossed with male mice carrying homozygous *Cx3cr1*-CreERT2, heterozygous floxed *Stim1* and *Stim2*, and homozygous PC::G5-tdT. Resulting offspring carrying *Cx3cr1*-CreERT2 were genotyped and classified as mutants (homozygous floxed *Stim1* and *Stim2*) or controls (heterozygous floxed *Stim1* and *Stim2*), along with intermediate genotypes. At 10 months of age, all mice received tamoxifen injections on five consecutive days (T1–T5) to induce Cre-mediated recombination in *Cx3cr1*-expressing microglia. Experiments commenced at 15 months of age. Diagram created with BioRender. (B) RT-PCR gel electrophoresis confirming the purity of isolated microglia. cDNA prepared from whole-brain RNA and from isolated microglia was amplified using primers for the housekeeping gene *Actb*, the microglial marker *P2ry12*, the neuronal marker *NeuN* (*Rbfox3*), the astrocyte marker *Gfap*, and the oligodendrocyte marker *Olig1*. Enrichment of *P2ry12* and reduced expression of non-microglial markers in the microglia fraction confirmed successful isolation. (C, D) Quantitative RT-PCR analysis of *Stim1* (C) and *Stim2* (D) mRNA in isolated microglia from control and mutant mice. Relative expression was calculated using the 2^−ΔΔCq^ method with *Actb* (β-actin) as the endogenous reference; each symbol is one animal and bars show mean ± SD of the plotted 2^−ΔΔCq^ values. *Stim1* mRNA was reduced by 66.9% and *Stim2* mRNA by 65.7% in mutant microglia relative to controls. n = 3 mice per group. *p < 0.05, unpaired one-tailed t-test on ΔCq values (*Stim1*, p = 0.041; *Stim2*, p = 0.014); the one-tailed test was specified in advance because excision of the floxed exons can only reduce the transcript.

*Calcium imaging* (Figure 2). Duration, peak amplitude (ΔF/F0), and area under the curve were measured for each microglial cell. Genotype is assigned at the animal, so the mouse was the unit of analysis. Each animal was reduced to one mean per parameter (n = 3 control, 4 mutant mice), and genotypes were compared with an exact permutation test of all 35 assignments of the seven mice to the two groups. The smallest p value this test can return is 1/35 = 0.029. As a supporting analysis, cell-level measurements were fit with a linear mixed-effects model, with genotype as a fixed effect and mouse as a random intercept. Inference used 5 residual degrees of freedom (mice − 2). Peak amplitude and AUC were log-transformed because cell-level residuals were right-skewed; duration was analyzed untransformed. Cells with no detectable response (duration, peak, and AUC all zero) were omitted from the magnitude comparisons, leaving 109 of 130 cells (44 of 48 control, 65 of 82 mutant). Permutation tests and mixed models were fit in Python (SciPy and statsmodels).

**Figure 2.**
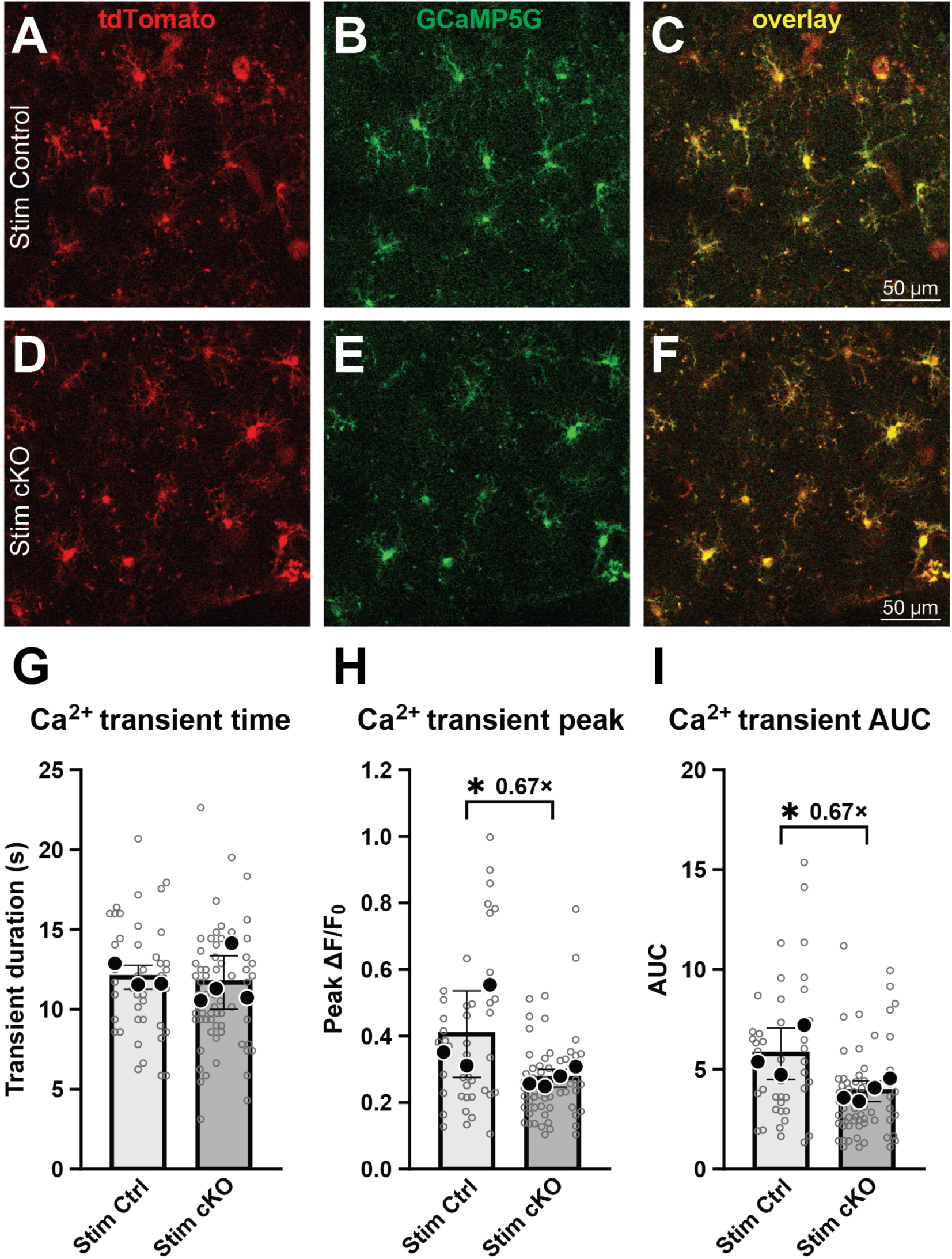
Microglia lacking *Stim1* and *Stim2* demonstrate reduced calcium transients during CSD. (A-F) Panels show representative 2p images of tdTomato (red) and GCaMP5G (green) signals, along with the signal overlay immediately after CSD depolarization passes. (A-C) Control animal; (D-F) a microglial *Stim* cKO. Scale bar, 50 µm. (G–I) Quantification of CSD-evoked calcium transients. (G) Transient duration, (H) peak amplitude (ΔF/F₀) and (I) area under the curve. Small circles show individual microglia, with a distinct horizontal position for each mouse so that cells from the same animal can be identified; large symbols show the mean for one mouse. The horizontal bar and whiskers show the mean ± SD of the per-mouse means. Statistical comparison used one mean per animal, with an exact permutation test over all 35 possible assignments of the seven mice; 0.029 is the smallest p-value this design can return. Peak amplitude was reduced by 32.7% and AUC by 32.5% in *Stim* cKO (both p = 0.029), while transient duration was unchanged (p = 0.74). n = 7 mice (3 controls, 4 Stim cKO) and 109 responding microglia of 130 tracked (44 of 48 control, 65 of 82 mutant). *p < 0.05; ns, not significant.

*MRI at 6 hours* (Figure 3). Eleven males were analyzed (6 control, 5 cKO). T2 infarct volume and the CBF measures, which were positive and right-skewed, were compared with a gamma generalized linear model (GLM) with a log link. Absolute lesion CBF was the pre-specified primary perfusion measure, and relative CBF (% of the contralateral region) is reported alongside it. Absolute CBF was chosen because ischemic viability thresholds are defined on the absolute scale, and because the ratio compounds estimation error from two separately measured regions. Sensitivity analyses of the lesion CBF comparison are listed in Table SM4. Relative T2 signal and the ADC measures were compared with unpaired Welch t-tests on the raw values. Effects are reported as cKO/control ratios of means (fold change) with 95% confidence intervals (Fieller’s method for the Welch tests).

**Figure 3.**
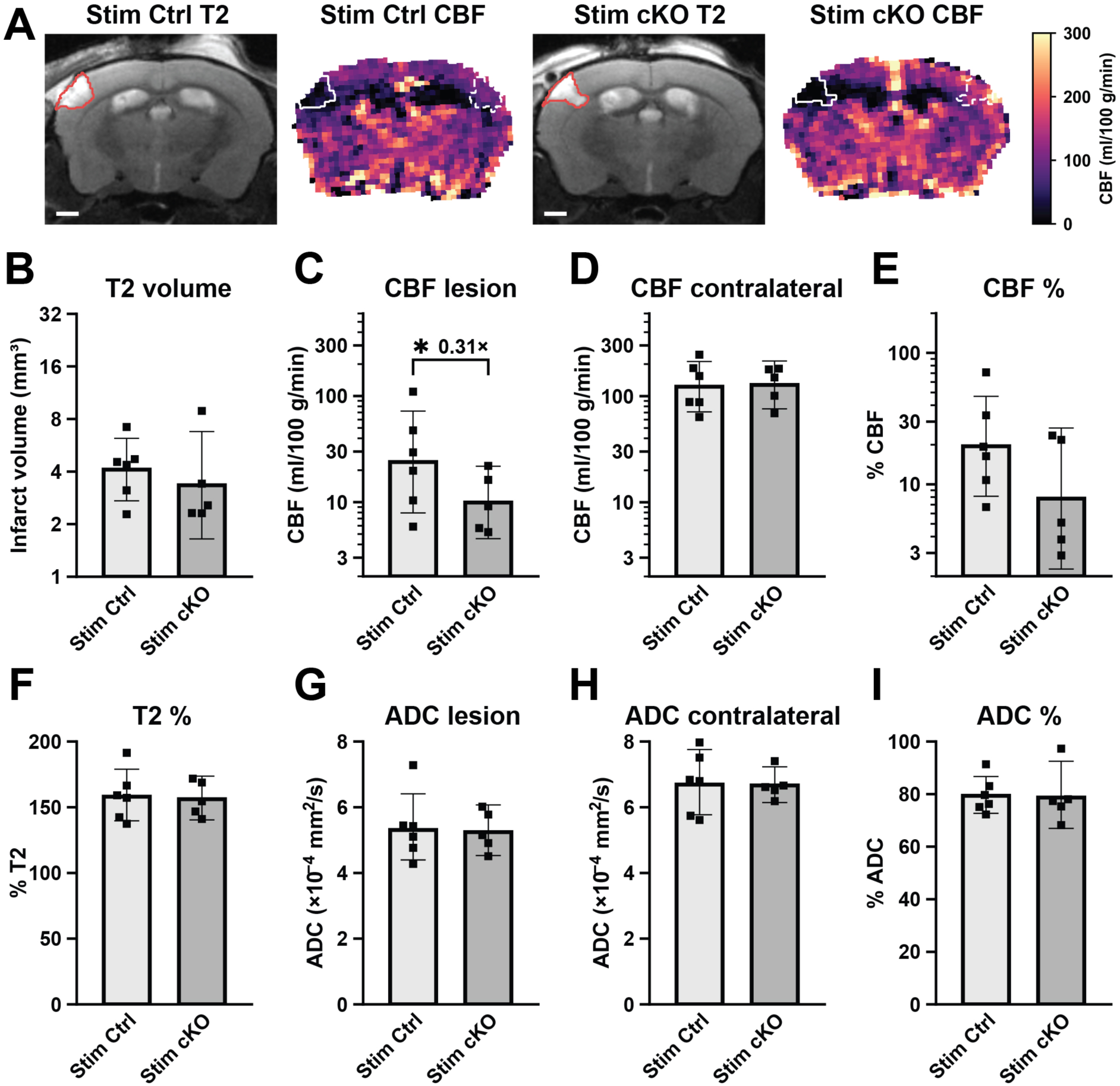
Six hours after dMCAo, infarct size, edema and diffusion are similar between genotypes, with lower lesion perfusion in *Stim* cKO mice. (A) Representative coronal T2-weighted images and cerebral blood flow (CBF) maps from a control (*Stim* Ctrl) and a mutant (*Stim* cKO) male, 6 h after occlusion of the right distal MCA. The two animals were matched for rostro-caudal level. Each T2 image is taken at the mid-plane of the 1-mm arterial spin labeling (ASL) slice, and the red outline marks the T2-defined infarct. On the CBF maps, the solid white line marks the same infarct mapped onto the ASL slab, and the dashed line marks its mirror image in the contralateral hemisphere, the within-animal reference region. Images follow radiological convention, with the right (stroke) hemisphere on the viewer’s left. CBF scale, 0– 300 ml/100 g/min. The maps show pixel CBF fitted on all 16 inversion times, as used in C–E. Scale bars, 1 mm. (B) T2-defined infarct volume (mm^3^). Each value is the mean of two independent tracings (two investigators, coronal plane). (C–E) CBF measured by flow-sensitive alternating inversion recovery (FAIR) ASL, fitted pixel by pixel on 16 inversion times and averaged over each region (Methods). (C) CBF within the infarct. (D) CBF in the mirrored contralateral region. (E) Lesion CBF as a percentage of the contralateral value in the same animal (% CBF). (F) Relative T2-weighted signal: mean signal within the infarct as a percentage of the mirrored contralateral region (% T2). Because the infarct is outlined as T2-hyperintense tissue, values exceed 100% by definition. (G–I) Apparent diffusion coefficient (ADC). (G) Mean ADC within the infarct. (H) Mean ADC in the mirrored contralateral region. (I) Lesion ADC as a percentage of the contralateral value (% ADC). The T2-defined masks were carried onto the diffusion images by registration. We excluded slices and diffusion directions corrupted by motion, and voxels with CSF-like diffusion (ADC > 1.5 × 10^−3^ mm^2^/s). Each symbol is one male mouse. n = 11 (6 control, 5 cKO) of 14 imaged. One mouse died during imaging, and two cKO mice were excluded before analysis because their infarct was at or below the T2 detection limit (∼0.5% of brain volume). Bars show the geometric mean, and error bars show its 95% confidence interval. Axes are logarithmic in B–E and linear in F–I. Lesion CBF was lower in *Stim* cKO (0.31-fold, p = 0.029) (C); sensitivity analyses are given in Table SM4. No other comparison was significant: infarct volume, 0.89-fold (p = 0.74) (B); contralateral CBF, 0.99-fold (p = 0.97) (D); % CBF, 0.43-fold (p = 0.13) (E); % T2, 0.98-fold (p = 0.81) (F); lesion ADC, 0.98-fold (p = 0.83) (G); contralateral ADC, 0.99-fold (p = 0.88) (H); and % ADC, 1.00-fold (p = 0.95) (I). Fold changes are cKO/control ratios of group means. p values are from a gamma GLM with a log link (B–E) or unpaired Welch t-tests (F–I). *p < 0.05; comparisons without a bracket were not significant.

*T2 infarct volume at 48 hours* (Figures 4B–D and S4B). T2 infarct volumes were positive, right-skewed, and showed variance that increased with the mean. Genotype effects were therefore estimated with a gamma generalized linear model and a log link, which yields the ratio of means directly. Effects are reported as fold-change with 95% confidence intervals. Sex was a pre-specified biological variable. Volumes were analyzed within each sex, and the genotype × sex interaction was tested with an interaction term in the gamma GLM (F test with estimated dispersion). The same direction and significance were obtained with Welch’s t-test, a t-test on log-transformed values, the Mann–Whitney U test, and a permutation test of group means.

**Figure 4.**
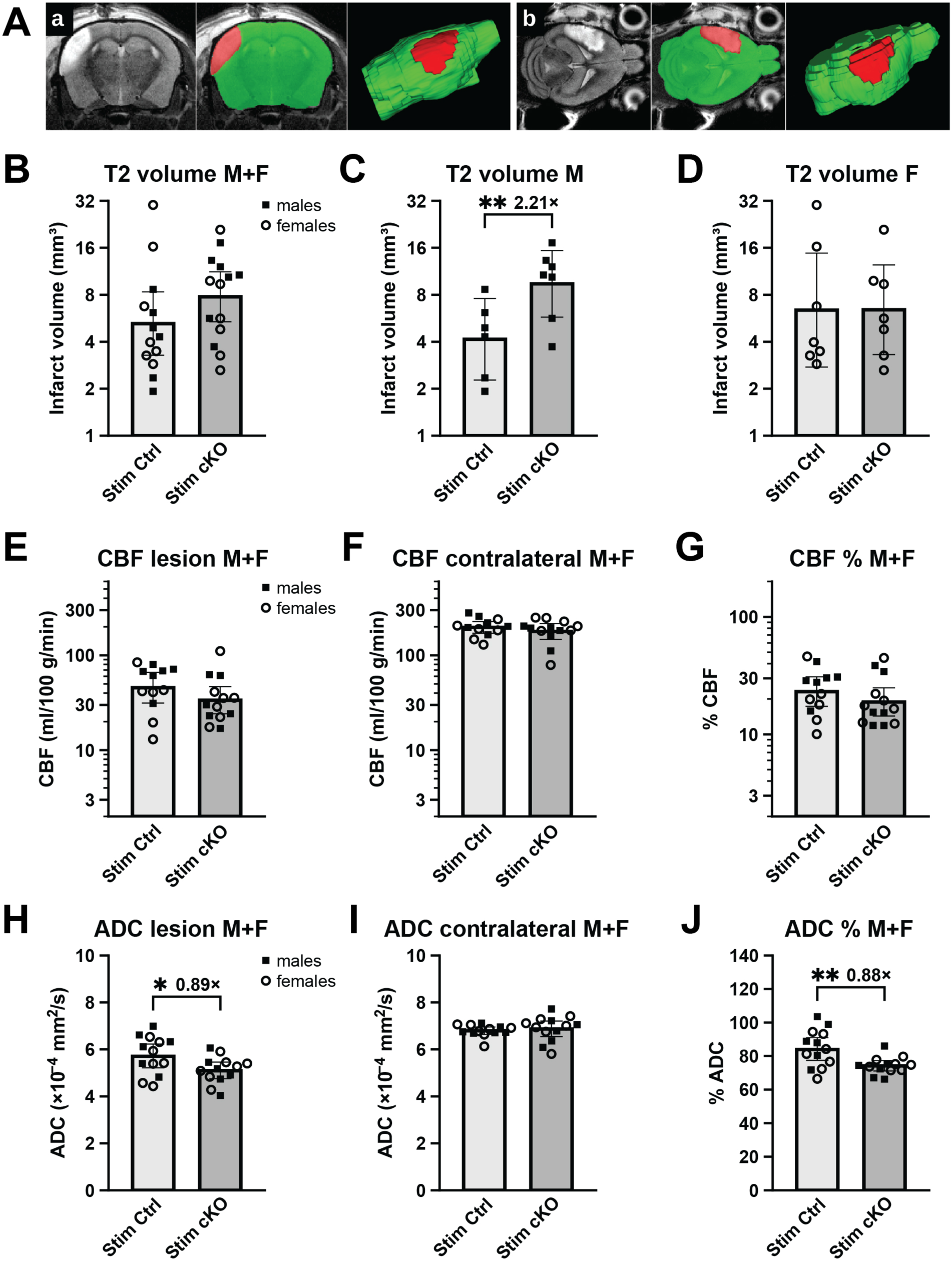
MRI-based quantification of infarct volume and stroke severity 48 hours after dMCAo. (A) Segmentation of the ischemic injury from T2-weighted MRI. The infarct (red) and whole brain (green) were traced in ITK-SNAP on every slice of the 48-hour T2 stacks. (a) Coronal series: unsegmented slice, the same slice with masks applied, and the 3D reconstruction from the coronal segmentation. (b) Axial series in the same order. Where the masks overlap, the infarct label is drawn on top. (B–D) T2-defined infarct volume (mm^3^) in control (*Stim* Ctrl) and mutant (*Stim* cKO) mice: both sexes pooled (B), males (C) and females (D). Each symbol is the mean of four tracings of that animal (two investigators × two planes). Infarct volume was larger in male mutants (2.21-fold, p = 0.0030) (C) but not in females (0.85-fold, p = 0.74) (D) or in the pooled cohort (1.27-fold, p = 0.47) (B). The genotype × sex interaction was not significant (p = 0.12). n = 27 (males: 6 control, 7 cKO; females: 7 per genotype). (E–G) Cerebral blood flow (CBF) measured by flow-sensitive alternating inversion recovery (FAIR) arterial spin labeling, fitted pixel by pixel on 13 inversion times and averaged over each region (Methods), both sexes pooled. (E) CBF within the T2-defined infarct. (F) CBF in the mirrored contralateral region. (G) Lesion CBF as a percentage of the contralateral value in the same animal (% CBF). None significantly differed by genotype: 0.76-fold (p = 0.22), 0.92-fold (p = 0.40) and 0.83-fold (p = 0.33), respectively. n = 25 (control: 6 males, 6 females; cKO: 6 males, 7 females). (H–J) Apparent diffusion coefficient (ADC), both sexes pooled. (H) Mean ADC within the T2-defined infarct. (I) Mean ADC in the mirrored contralateral region. (J) Lesion ADC as a percentage of the contralateral value (% ADC). Lesion ADC was lower in *Stim* cKO (0.89-fold, p = 0.033), and so was % ADC (0.88-fold, p = 0.0078); contralateral ADC did not differ (1.01-fold, p = 0.59). n = 26 (each genotype: 6 males, 7 females). Masks and voxel exclusions were as in Figure 3G–I. Throughout, each symbol is one animal (squares, males; circles, females). Bars show the geometric mean, and error bars show its 95% confidence interval. Axes are logarithmic in B–G and linear in H–J. Fold changes are cKO/control ratios of group means. P-values are from a gamma GLM with a log link (B–G) or unpaired Welch t-tests (H–J). Of 34 mice subjected to dMCAo, seven were excluded before analysis because the infarct was at or below the T2 detection limit (∼0.5% of brain volume), judged at segmentation and blind to genotype. Circulation failed in two of the remaining 27 mice during the imaging session, as shown by absent perfusion on ASL. One cKO male had already lost circulation by the DWI scan (whole-brain ADC 35% below the cohort median) and was excluded from E–J. One control female lost circulation between the DWI and ASL scans and was excluded from E– G only. *p < 0.05, **p < 0.01; comparisons without a bracket were not significant.

*Infarct growth from 6 to 48 hours* (Figure S4C). Because the 6-h cohort was male-only, growth was tested in males (6 h: 6 control, 5 cKO; 48 h: 6 control, 7 cKO) with gamma GLMs (log link). Growth within each genotype compared the two time points (fold change = 48-h/6-h ratio of means). Whether growth differed between genotypes was tested with the time × genotype interaction in a joint model containing all four groups. Sensitivity analyses used a single investigator’s coronal tracings at both time points, infarct volume as a percentage of brain volume, and 48-h volumes adjusted for hemispheric swelling.

*CBF at 48 hours* (Figure 4E–G). Lesion, contralateral and relative CBF were compared with a gamma GLM (log link); n = 25 (12 control, 13 cKO). Sex and the genotype × sex interaction were tested in the same model. The association between lesion CBF and lesion ADC across animals was assessed with Spearman’s correlation and, to account for genotype, with a linear model of lesion ADC on log lesion CBF and genotype.

*MRA analysis* (Figure S5). Vessel length density was compared between the stroke side and the contralateral side within each animal using the two-sided Wilcoxon signed-rank test, pooled across all 25 animals (control male 6, control female 6, cKO male 6, cKO female 7).

Group sizes of six preclude per-group testing because the smallest attainable two-sided Wilcoxon p is 0.031 and requires unanimity, so the four groups are displayed but not tested individually. Sex and genotype were compared with two-sided Mann–Whitney U tests; four-group comparisons used Kruskal–Wallis. Related tests were Holm–Bonferroni-adjusted within family: the seven hemispheric measures in (A), and the four ROI × side combinations in (E) and (F), with genotype and sex treated as separate families. Adjusted p-values are given where the correction applies. Hemispheric ratios and ROI volumes are mean ± SD; all other data are mean ± SEM. Analysis was performed in Python 3.13.13 (numpy 2.3.5, scipy 1.15.3, pandas 3.0.3, SimpleITK 2.5.5, scikit-image 0.26.0, ANTsPy 0.6.3 and matplotlib 3.10.9). Brains were registered to the Perens stereotaxic MRI mouse atlas (brainglobe-atlasapi 2.3.1, perens_stereotaxic_mri_mouse_25um).

*ADC measures* (Figure 4H–J). Lesion ADC, contralateral ADC and relative ADC were compared on the raw scale with unpaired Welch t-tests, because group variances were unequal (Levene p = 0.009 for relative ADC); n = 26 (13 per genotype). The primary analysis pooled both sexes. Sex-stratified Welch tests are reported in the Results, and the genotype × sex interaction was tested with a linear model.

*Hemispheric swelling* (Figure S4A). Swelling across the cohort was tested with a paired t-test of each animal’s infarcted-slab asymmetry against its lesion-free-slice asymmetry, and genotypes were compared with Welch’s t-test (n = 27). The same paired test was applied to the 11 males imaged at 6 h.

*Segmentation and inclusion.* Each 48-hour T2 infarct was traced independently by two investigators in the coronal and axial planes, and the four measurements were averaged (intraclass correlation = 0.99 for the four-measurement mean and 0.98 for a single tracing).

Animals in which distal MCA occlusion produced a lesion at or below the reliable detection limit of the T2 sequence (∼0.5% of total brain volume) were identified during segmentation, blind to genotype, and excluded before analysis. This criterion removed 7 of 34 mice (4 controls, 3 mutants). Repeating the male volume analysis after restoring every measurable infarct did not change the result (2.19-fold; 95% CI 1.24–3.87; p = 0.0072). No animal was excluded as a statistical outlier. Of 14 males imaged at 6 h, one died during imaging, and two cKO mice with infarcts at or below the T2 detection limit were excluded by the same criterion. At 48 h, circulation failed in two of the 27 included mice during the imaging session, as shown by absent perfusion on ASL. One of them, whose normal-tissue ADC was already 35% below the cohort median at the DWI scan, was excluded from the CBF and ADC analyses; the other was excluded from the CBF analysis only.

The data that support the findings of this study are available from the corresponding authors upon reasonable request.

## RESULTS

### Genetic deletion of *Stim* genes in microglia

We sought to ablate CRAC channel function by deleting the genes encoding the ER calcium sensors *Stim1* and *Stim2*. In immune cells, *Stim1* is generally expressed at higher levels than *Stim2* (Korshunov & Prakriya, 2025; Kraft, 2015). However, we combined floxed alleles of both *Stim* genes to ensure complete disruption of CRAC function. To achieve conditional *Stim* knockout (cKO) specifically in microglia, we used the microglia-selective *Cx3cr1-CreERT2* driver (Yona et al., 2013). We have also included the PC::G5-tdT reporter cassette, which expresses the green calcium indicator GCaMP5G and the red fluorescent reporter tdTomato (Gee et al., 2014). Experimental animals of both sexes carrying the desired genotype were induced at 10 months of age using five consecutive tamoxifen injections. The experiments were conducted 5 months after induction (Figure 1A). It is worth noting that *Cx3cr1-CreERT2* also recombines circulating CX3CR1+ monocytes and other non-parenchymal myeloid cells at the time of tamoxifen administration. However, the circulating cells are replaced by unrecombined bone-marrow progeny within weeks (Goldmann et al., 2013; Yona et al., 2013). Thus, the 5-month post-induction interval leaves the *Stim* deletion only in long-lived CX3CR1+ populations, primarily in parenchymal microglia, and also in a subset of self-renewing border-associated and meningeal macrophages (Goldmann et al., 2016).

Deletion of both *Stim* genes was confirmed by RT-PCR at the time of experimentation. Microglia were isolated from the brains of three male control and three male cKO mice using CD11b MicroBeads, and isolate purity was verified with markers for major brain cell types (Figure 1B). RT-PCR was performed with *Stim1* and *Stim2* primers, using *Actb* as the reference gene (Table 1). Because Cre-mediated excision of the floxed exons can only lower the transcript, expression was compared on the ΔCq scale with a one-tailed t-test. *Stim1* mRNA fell by 66.9% in mutant microglia (ΔΔCq = 1.60 cycles; p = 0.041), and *Stim2* mRNA by 65.7% (ΔΔCq = 1.55 cycles; p = 0.014) (Figures 1C and 1D). Both reductions confirm efficient disruption of *Stim* transcripts in microglia. The functional consequence is demonstrated in the calcium imaging experiment that follows.

### *Stim* cKO microglia exhibit reduced calcium influx during spreading depolarizations

We next examined how *Stim* gene deletion affects microglial calcium influx using in vivo two-photon imaging through cranial windows. We previously identified microglial calcium loading during peri-infarct depolarizations following ischemic injury (Liu et al., 2021). However, the stochastic nature of these peri-infarct depolarizations poses a significant experimental challenge. To obtain sufficient reproducible measurements, we induced cortical spreading depolarizations with KCl. The KCl microinjections were delivered through burr holes in the visual cortex, and the responses were recorded in the somatosensory cortex area (Figure S1A). Each stimulus elicited a calcium wave in the imaging field with an approximately 60-second delay.

Microglial responses were captured using the structural reporter tdTomato and the calcium indicator GCaMP5G (Figure 2A-F). A total of 12 mice were used in the experiments, and each animal underwent 3-6 KCl-induced depolarizations. After removing recordings with motion artifacts, seven mice (three controls and four *Stim* cKO mice) were used for final analysis. Microglial somas were segmented using the tdTomato signal, and the GCaMP signal was calculated within each segmented region (Figure S1). In each field of view, 3-6 cells were suitable for analysis. Fluorescence intensity traces were smoothed, and the duration, peak amplitude (ΔF/F_0_), and area under the curve (AUC) were calculated (Figure S1).

Intracellular calcium activity was reduced in *Stim* cKO microglia. Because each mouse contributed several cells, we compared genotypes using one mean per animal, and assessed significance with an exact permutation test over all 35 possible assignments of the seven mice. Mean AUC was 32.5% lower in *Stim* cKO (5.772 ± 1.291 to 3.896 ± 0.517, p = 0.029), and mean peak amplitude was 32.7% lower (0.406 ± 0.130 to 0.273 ± 0.027, p = 0.029) (Figure 2). Ranked by mean per animal, the three control mice held the three highest values of the seven for both measures, the most extreme of the 35 possible groupings. A linear mixed-effects model fitted to all 109 responding cells, with mouse as a random intercept, gave the same conclusion for AUC (p = 0.016). Notably, the duration of calcium transients remained unchanged (12.0 ± 0.8 s versus 11.7 ± 1.7 s, p = 0.74).

Of 130 tracked microglia, 109 showed a detectable calcium response and were included in these measurements. Non-responding cells were somewhat more frequent in mutants (17 of 82, 21%) than in controls (4 of 48, 8%), although this difference did not reach significance at the animal level (p = 0.18). Together, these results indicate that *Stim* gene deletion attenuates microglial calcium responses to CSD by approximately one third.

### Assessing the effects of decreased calcium signaling in the context of ischemic injury

Having confirmed that *Stim* gene deletion decreases microglial calcium signaling, we next asked whether this reduction would translate into improved outcomes following ischemic stroke.

We reasoned that diminished calcium load in microglia would reduce proinflammatory polarization and, consequently, mitigate ischemic injury. To test this prediction, we initially used the well-established model of proximal middle cerebral artery occlusion (pMCAo) (Norat et al., 2023; Tvrdik et al., 2019). We first prioritized pMCAo because it occludes the MCA at its origin, and the resulting ischemic region spans both cortical and subcortical structures, potentially offering a more robust approach for detecting genotype-dependent differences. In this initial cohort, we subjected three controls and three cKO mice to pMCAo and assessed stroke volume 48 hours later using a 9.4T MRI scanner with a T2-weighted 3D sequence (Figure S2). Manual segmentation revealed an average stroke volume of 101.8 ± 28.94 mm³ in controls and 130 ± 21.67 mm³ in cKO mice (Figure S2E), corresponding to an increase from 21.4% to 25.7% of total brain volume (Figure S2F), suggesting a trend toward larger infarcts in the cKO group. However, the extensive tissue damage induced by pMCAo resulted in high mortality and large, variable infarct volumes, which together limited statistical power and precluded definitive conclusions.

To address these limitations, we adopted the distal MCAo (dMCAo) model, which confines ischemic injury to the MCA cortical territory and substantially reduces both mortality and outcome variability. Importantly, the resulting lesion is restricted to the somatosensory cortex, the same region in which we characterized microglial calcium responses, thereby allowing a more direct correlation between microglial physiology and stroke outcome.

### Six hours after dMCAo, lesion perfusion tends to be lower in *Stim* cKO mice despite similar infarct size, edema, and diffusion

To limit ischemic injury to the somatosensory cortex, we employed distal middle cerebral artery occlusion (dMCAo), as previously described (Norat et al., 2023) and depicted in Figure S3.

Initially, we examined the effects of microglial *Stim* mutations during the hyperacute stage, 6 h after ischemic stroke. At this stage, salvageable areas still surround the ischemic core, forming the penumbra. To distinguish the core and penumbra, we combined several MRI methods to characterize these ischemic regions. We optimized T2, diffusion-weighted imaging (DWI), and arterial spin labeling (ASL) approaches, which are sensitive to edema, cell death, and tissue blood perfusion, respectively. We treated and imaged 14 mice and analyzed cerebral ischemia in 11 male mice (6 controls, 5 cKO) six hours after occlusion (Figure 3).

We first examined the 6-h infarcts with T2-weighted imaging. Infarct volume, determined by manual segmentation, did not differ between genotypes (4.37 ± 1.68 mm^3^ in controls and 3.90 ± 2.83 mm^3^ in *Stim* cKO; 0.89-fold, p = 0.74) (Figure 3B). The T2 signal inside the infarct was also similar. Relative to the mirrored contralateral region, it reached (159 ± 19)% in controls and (157 ± 13)% in *Stim* cKO (0.98-fold, p = 0.81) (Figure 3F). At 6 h, the two genotypes therefore carried infarcts of the same size and with comparable edema.

Lesions of equal size can still differ in severity. Cell swelling, membrane integrity and tissue water mobility all vary, and the apparent diffusion coefficient (ADC) reports them quantitatively. Perfusion can differ as well. We therefore measured ADC and cerebral blood flow (CBF) within the same T2-defined lesions. Each value was referenced to the mirror image of the lesion in the contralateral hemisphere of the same animal (Figure 3A; Figure S4D).

Diffusion was restricted to the same extent in both genotypes. Lesion ADC was (5.38 ± 1.03) × 10^−4^ mm^2^/s in controls and (5.27 ± 0.62) × 10^−4^ mm^2^/s in *Stim* cKO (0.98-fold, p = 0.83) (Figure 3G). Contralateral ADC did not differ either ((6.74 ± 0.94) × 10^−4^ mm^2^/s in controls and (6.68 ± 0.45) × 10^−4^ mm^2^/s in *Stim* cKO; 0.99-fold, p = 0.88) (Figure 3H). Relative ADC was (80 ± 7)% and (79 ± 11)%, respectively (1.00-fold, p = 0.95) (Figure 3I). The cytotoxic component of the injury was thus already present at 6 h, and it was equal in the two genotypes.

Lesion CBF was also reduced below the contralateral level in both genotypes. In contrast to ADC, however, the lesion side CBF was lower in *Stim* cKO mice (11.6 ± 7.3 versus 37.1 ± 38.6 ml/100 g/min; 0.31-fold, 95% CI 0.11–0.89, p = 0.029) (Figure 3C), whereas contralateral CBF did not differ (136 ± 50 versus 137 ± 70 ml/100 g/min; 0.99-fold, p = 0.97) (Figure 3D). The difference was therefore confined to the lesion and did not reflect a global change in flow. Expressed as a percentage of the contralateral value, lesion CBF followed the same trend but did not reach significance ((11.4 ± 10.3)% versus (26.3 ± 23.8)%; 0.43-fold, p = 0.13) (Figure 3E).

These findings require cautious interpretation because lesion CBF varied widely among controls. One of the six control lesions retained high flow (110 ml/100 g/min), whereas all five *Stim* cKO lesions stayed below 22 ml/100 g/min. Without that control, the difference was smaller and not significant (0.52-fold, p = 0.12). Tests that do not assume a gamma distribution gave p = 0.12–0.17, and the comparison weakened when only the 13 inversion times shared with the 48-h protocol were used (0.38-fold, p = 0.064) (Table SM4). We therefore regard it as a signal rather than a settled difference. Nevertheless, three of the six control infarcts retained flow above 25 ml/100 g/min, and no *Stim* cKO infarct did.

Counterintuitively, lesion perfusion was lower in Stim cKO mice, but lesion size and injury severity did not differ from controls. One possible explanation is timing. ADC decreases within minutes of ischemia, while T2 hyperintensity builds up over the following hours (Gerriets et al., 2004; Gerriets et al., 2009; Moseley et al., 1990). By 6 h, injury had reached a similar extent in both genotypes. However, a deeper perfusion deficit may affect penumbral tissue that is still viable, with consequences that would only become evident as the infarct matures. If so, *Stim* cKO mice should develop larger or more severely injured infarcts at later stages. We tested this prediction 48 h after occlusion.

### At 48 h post-dMCAo, *Stim* cKO males have larger infarcts, and *Stim* cKO lesions show lower ADC

We imaged a second, independent dMCAo cohort 48 hours after occlusion. Thirty-four mice of both sexes were scanned with the same T2-weighted, diffusion-weighted and perfusion-weighted sequences, together with MR angiography.

First, we examined hemispheric swelling within the infarcted region. At 48 h, the ipsilateral hemisphere was enlarged by (1.4 ± 1.8)% (20 of 27 mice; p = 0.009). This excess volume was 1.7 mm³, 22 % of the T2 lesion volume. Swelling did not differ between genotypes (*Stim* Ctrl 1.7 %, *Stim* cKO 1.0 %; p = 0.35) (Figure S4A). Notably, no swelling was detectable at 6 h ((−1.9 ± 2.8)%; p = 0.16).

The 48-hour T2 series were acquired in the coronal and axial planes and segmented in ITK-SNAP (Figure 4A). Each infarct volume is the mean of four independent tracings (two investigators × two planes). Sex was a pre-specified biological variable; males and females are shown grouped and separately in Figure 4 B-D (males: 6 control, 7 mutant; females: 7 per genotype).

In males, T2 infarct volume was 2.21-fold larger in Stim cKO than in controls (4.72 ± 2.50 versus 10.44 ± 4.56 mm^3^; geometric means 4.15 and 9.40 mm^3^; 95% CI 1.31–3.74, p = 0.0030) (Figure 4C). The same difference was present when the infarct was expressed as a percentage of brain volume ((1.00 ± 0.55)% versus (2.22 ± 1.03)%; 2.21-fold; 95% CI 1.28–3.83, p = 0.0046) (Figure S4B).

To examine how injury evolved over time, we compared the 48-h males with the 6-h cohort, which was all male. Because the two time points came from independent cohorts, we compared the two time points within each genotype and tested whether infarct growth differed between genotypes with a time × genotype interaction (gamma GLMs). In controls, infarct volume did not change between 6 h and 48 h (4.37 ± 1.68 versus 4.72 ± 2.50 mm3; 1.08-fold, 95% CI 0.64–1.82, p = 0.78). In *Stim* cKO mice, it increased 2.68-fold (3.90 ± 2.83 versus 10.44 ± 4.56 mm^3^; 95% CI 1.39–5.15, p = 0.0032). Importantly, this growth differed significantly between genotypes (time × genotype interaction, 2.48-fold; 95% CI 1.07–5.73, p = 0.033) (Figure S4C). Thus, control infarcts were essentially complete at 6 h, whereas mutant infarcts continued to expand over the following two days.

Importantly, the result did not depend on how the infarcts were measured. The volumes in Figures 3 and 4 combine different tracings by several raters, so we repeated the analysis with the single investigator who traced the coronal series at both time points (Figure S4C). Mutant infarct increase (2.57-fold, p = 0.0047) and the interaction (2.52-fold, p = 0.028) were unchanged. The same was true for infarct volume expressed as a percentage of total brain volume (interaction 2.48-fold, p = 0.036). Edema did not explain the infarct growth either. After subtracting the swelling-related excess volume at 48 h (Figure S4A), mutant infarcts were still 2.16-fold larger than at 6 h (p = 0.029), whereas control infarcts did not grow (0.70-fold, p = 0.33; interaction p = 0.026). Together, these data point to the hours and days after occlusion as the critical window of vulnerability for the *Stim* cKO phenotype, when penumbral tissue is either salvaged or recruited into the infarct.

In 48-hour dMCAo females, infarct volume did not differ by genotype, either in absolute terms (0.85-fold; 95% CI 0.32–2.26, p = 0.74) or as a percentage of brain volume (0.86-fold, p = 0.77). With seven animals per group, this comparison had 39% power to detect the effect size seen in males and would have required a 3.8-fold difference for 80% power. The genotype × sex interaction was 2.62-fold (95% CI 0.78–8.74) and was not significant (p = 0.12), so the cohort does not establish that the effect is sex-specific. The pooled analysis across both sexes was also not significant (1.27-fold, p = 0.47). Returning every animal excluded for a sub-threshold infarct left the male result unchanged (2.19-fold; 95% CI 1.24–3.87, p = 0.0072).

To rule out baseline differences in vascular anatomy, we imaged the cerebral arteries with time-of-flight MR angiography during the same 48 h imaging session (Figure S5). Usable angiograms were obtained in 25 of the 27 animals; two were excluded for insufficient arterial contrast. The circle of Willis was complete, and both internal carotid arteries were patent in every animal. Hemispheric angioarchitecture was symmetric. The right-to-left ratio of total vessel length was 1.010 ± 0.124, not different from unity (p = 0.58). Notably, it did not differ by genotype or sex (all Holm-adjusted p > 0.05; four-group Kruskal–Wallis, p = 0.39) (Figure S5A). The occlusion itself was also uniform; vessel length density within the stroke territory decreased from 0.291 ± 0.019 in the contralateral region to 0.124 ± 0.021 mm per mm^3^ of ROI on the stroke side, reduced in 21 of 25 animals (p = 4.2 × 10⁻⁶) (Figure S5E). An unaffected control territory shifted 4 mm caudally in the visual cortex showed no such difference (p = 0.41) (Figure S5F). The deficit is therefore focal, not hemispheric. Importantly, neither measure differed by genotype or sex (all Holm-adjusted p ≥ 0.84). Together, these data argue against differences in large-artery anatomy or in the extent of occlusion as the cause of the larger infarcts in cKO males. The arterial tree was comparable across groups, and electrocoagulation removed a comparable amount of vessel in every animal. Thus, the phenotype distinguishing male *Stim* cKO mice is more likely attributable to tissue-intrinsic factors than to differences in large-vessel anatomy.

Perfusion within the infarct remained low at 48 h (Figure 4E-G). As with the 6 h studies, absolute lesion CBF was the primary measure. Lesion CBF averaged 51.9 ± 23.6 ml/100 g/min in controls and 39.2 ± 26.0 ml/100 g/min in *Stim* cKO mice, compared with contralateral values of 201 ± 44 and 186 ± 48 ml/100 g/min, respectively. As at 6 h, lesion flow was lower in cKO mice on average, but the difference was not significant in absolute terms (0.76-fold, p = 0.22) or relative to the contralateral region ((25.3 ± 11.0)% in controls versus (21.0 ± 11.0)% in cKO; 0.83-fold, p = 0.33). Contralateral CBF did not differ between genotypes (0.92-fold, p = 0.40).

None of these three measures differed between males and females (all p ≥ 0.37), and there was no genotype × sex interaction (all p ≥ 0.10).

This result was expected. By 48 h, tissue at risk has largely been either salvaged or recruited into the infarct, so residual flow within a completed infarct is a weaker readout of its fate. Flow nonetheless correlated with tissue state across animals; lesions with lower CBF had lower ADC (Spearman ρ = 0.71, p = 7.4 × 10^−5^), independent of genotype (p = 0.0002 for CBF in a linear model with genotype). We therefore turned to ADC, which reports tissue condition directly.

We examined the severity of tissue injury in 48 h Stim cKO and controls by determining ADC values in T2-positive lesions. We found that mean ADC within the T2-defined masks was significantly lower in *Stim* cKO than in controls (lesion ADC (5.76 ± 0.82) × 10^−4^ mm^2^/s in controls and (5.12 ± 0.58) × 10^−4^ mm^2^/s in Stim cKO; 0.89-fold, p = 0.033) (Figure 4H).

Contralateral ADC, obtained by mirror-transposing the T2 masks onto the opposite hemisphere (Figure S4D), did not differ between groups ((6.80 ± 0.26) and (6.89 ± 0.54) × 10^−4^ mm^2^/s, respectively; 1.01-fold, p = 0.59) (Figure 4I). Relative ADC (lesion/contralateral) was likewise reduced in the cKO mice ((84.6 ± 11.2)% in controls versus (74.3 ± 5.1)% in cKO; 0.88-fold, p = 0.0078) (Figure 4J). Specifically, relative ADC was lower in male *Stim* cKO ((88.5 ± 11.7)% in control males versus (73.6 ± 7.2)% in cKO males; 0.83-fold, p = 0.027) but not in females ((81.2 ± 10.4)% versus (74.9 ± 3.0)%; 0.92-fold, p = 0.17); the genotype × sex interaction was not significant (p = 0.22).

Interestingly, relative ADC moved in opposite directions in the two genotypes between 6 h and 48 h. At 6 h, it was the same in control and cKO males (79.6% and 79.2%; Figure 3I). By 48 h, it had risen to 88.5% in control males and fallen to 73.6% in cKO males. Neither change was significant on its own (controls p = 0.14; cKO p = 0.36). Notably, its direction matches infarct growth over the same interval, which was absent in controls and 2.68-fold in cKO males (Figure S4C). Together, these findings indicate that ischemic injury is more severe in *Stim* cKO mice, particularly males, than in mice with preserved microglial *Stim* expression.

After MRI, brains were stained with TTC. Quantifiable staining was obtained in 8 of 26 processed brains (2 control and 6 mutant males); no statistical test was performed. Mean TTC infarct volume was 1.22 mm^3^ in controls and 3.12 mm^3^ in cKO males, in the same direction as the T2 measurements (Figure S6). Taken together, loss of microglial *Stim1* and *Stim2* was associated with larger infarcts and greater diffusion restriction in male mice. A working model is shown in Figure 5.

**Figure 5.**
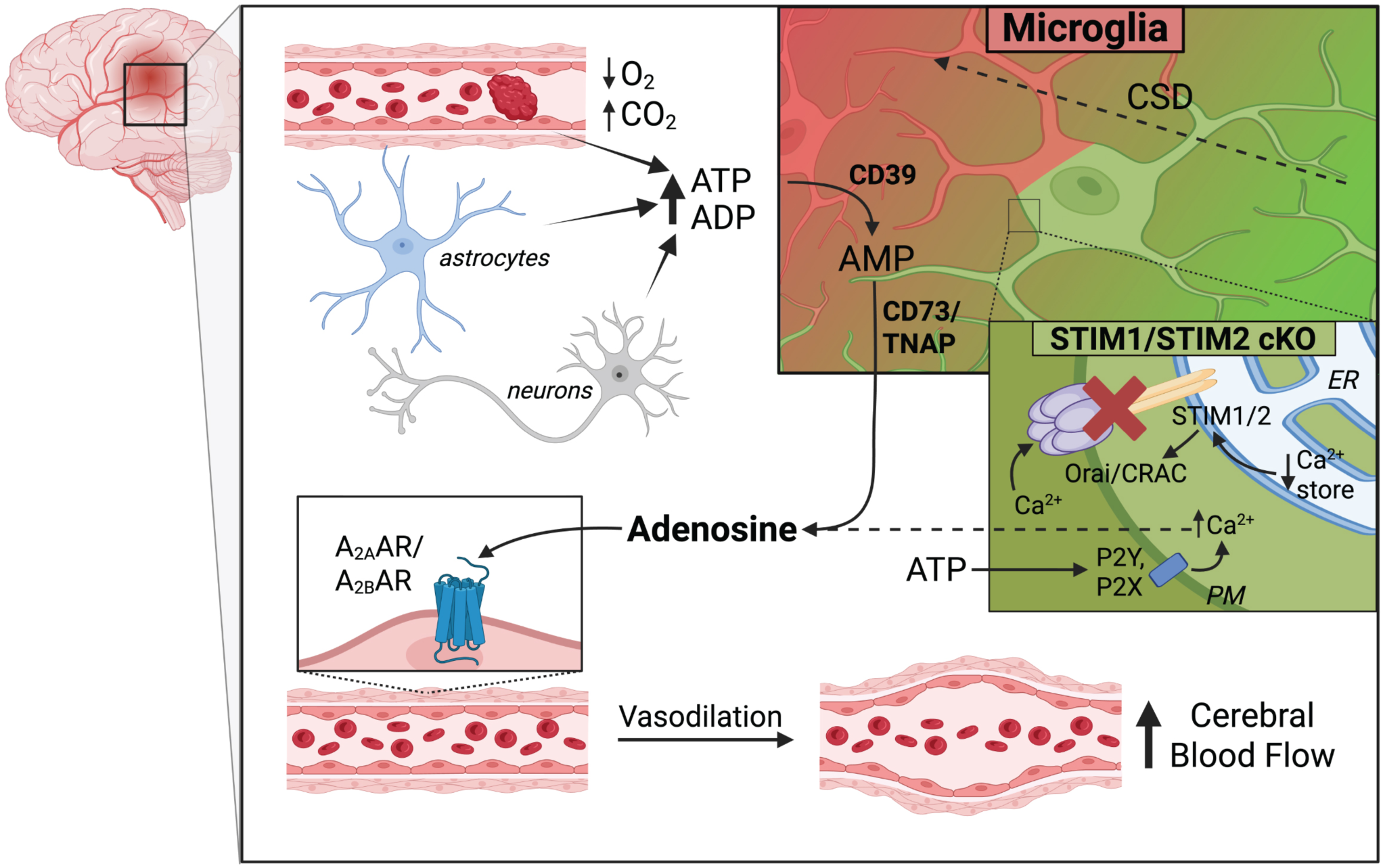
Proposed model: microglial CRAC channel signaling regulates the production of vasodilatory adenosine in the ischemic cortex. The schematic illustrates a cortical neurovascular unit within the peri-infarct zone. Arterial occlusion decreases tissue O₂ and increases CO₂, while recurrent cortical spreading depolarizations (CSDs) propagate through the adjacent cortex, indicated by the green-to-red gradient. Neurons, astrocytes, and endothelial cells release ATP and ADP (Csaszar et al., 2022). Microglia detect these nucleotides via P2Y receptors, primarily the high-affinity P2Y12 receptor, and through ionotropic P2X channels (Csaszar et al., 2022; Fu et al., 2025). Gq-coupled P2Y signaling depletes endoplasmic reticulum calcium stores; subsequently, Stim1 and Stim2 activate Orai/CRAC channels to maintain Ca²⁺ influx. Inhibition of this process reduces CSD-evoked microglial calcium transients (Liu et al., 2021), and this is the step disrupted in the microglial Stim1/Stim2 mutants examined in this study (red cross). Calcium-dependent motility facilitates P2Y12R-dependent perivascular filopodia that interact with the vessel wall (Csaszar et al., 2022). Microglial CD39 hydrolyzes ATP to ADP and AMP; CD73, or Tissue-Nonspecific Alkaline Phosphatase (TNAP) completes the conversion to adenosine, which then acts on perivascular A₂ₐ and, with lower affinity, A₂ᵦ receptors on smooth muscle cells and pericytes to increase local cerebral blood flow (Fu et al., 2025). Solid arrows represent steps demonstrated in the cited studies, while dashed arrows indicate steps inferred for the ischemic context. Notably, microglial adenosine production was demonstrated under hypercapnia rather than hypoxia (Csaszar et al., 2022). This pathway was not directly tested in the present study. Diagram created with BioRender.

## DISCUSSION

We have shown that microglial deletion of *Stim1* and *Stim2* reduces CSD-evoked calcium transients by approximately one-third, but worsens the subsequent ischemic injury. The key finding was a divergence in infarct evolution. The mice had comparable ischemic injury 6 h after occlusion, but by 48 h the cKO infarcts were 2.21-fold larger. Lesion perfusion was already lower in cKO mice at 6 h, before any detectable differences in infarct volume or diffusion, although this difference did not survive every sensitivity analysis. We propose that microglial STIM-mediated calcium signaling helps protect ischemic tissue as the infarct evolves (Figure 5).

To assess if baseline angiography is affected by microglial *Stim* ablation, we examined dMCAo mice with MRA. It showed no group differences in arterial anatomy, circle-of-Willis integrity, hemispheric vessel density, or the angiographic extent of vessel loss after electrocoagulation (Figure S5). These findings argue against the initial occlusion or gross arterial anatomy as the explanation for the group differences. Instead, they point to impaired tissue-level adaptation to ischemia, possibly involving collateral and microvascular perfusion.

Temporal analysis of infarct growth revealed that the differences between cKO and control phenotypes arise after the initial lesion has formed. Across the independent 6 h and 48 h cohorts, control infarct volumes did not change. Conversely, cKO infarct volumes increased 2.68-fold. The time × genotype interaction was significant (Figure S4C), showing that infarcts indeed expanded more in cKO mice than in controls. It remained significant with alternative tracing and normalization methods and after correction for swelling. The minor hemispheric swelling seen in most 48 h dMCAo mice did not differ by genotype, so edema is unlikely to explain the difference. In addition, we show lower ADC in 48 h cKO mice, further corroborating more severe tissue damage in the lesions lacking microglial *Stim*.

Our early perfusion measurements support this interpretation. Absolute lesion CBF was lower in 6 h cKO males than in controls (11.6 ± 7.3 versus 37.1 ± 38.6 ml/100 g/min; 0.31-fold, p = 0.029), while CBF in the mirrored contralateral region did not differ (p = 0.97). This was the earliest *Stim* genotype difference we detected in the occluded areas, while lesion volume and ADC remained comparable. The limited cohort size and dispersed control values warrant caution. The difference depended on one control with high residual flow and on the statistical test (Table SM4). Even so, the absolute CBF values suggest that perfusion differed before the infarct burden diverged.

This microglia-dependent temporal pattern seen in this study is consistent with the work of Császár et al. (Csaszar et al., 2022). The authors linked microglial calcium signaling to cerebrovascular regulation using the same CX3CR1-driven GCaMP5G–tdTomato reporter line. They showed that hypercapnia induced rapid microglial calcium changes, formation of P2Y12R-mediated perivascular contacts, and localized adenosine release. In contrast, microglial depletion impaired hypercapnia-induced vasodilation. Further, removal of microglia or P2Y12R blockade substantially impaired neurovascular coupling and cerebrovascular adaptation to cortical hypoperfusion after common carotid artery occlusion. These findings provide a plausible framework in which reduced STIM-dependent calcium signaling limits microglial support for vascular compensation. Our 6-hour CBF result fits into this mechanistic interpretation; a perfusion difference was already detectable before the infarct volumes diverged. The subsequent expansion in cKO males is therefore consistent with a failure to sustain perfusion sufficiently to preserve ischemic penumbra. However, Császár et al. studied the vascular responses under different experimental conditions, and their dependence on Stim1/Stim2 during focal ischemia remains to be demonstrated.

Further insight into how microglia regulate cerebral blood flow through adenosine production comes from the work of Fu et al. (2025), who demonstrated that the microglial ectonucleotidase CD39 is essential for mediating cerebrovascular reactivity (Fu et al., 2025). CD39 initiates the extracellular dephosphorylation cascade that converts ATP to ADP and AMP, eventually leading to adenosine generation and vasodilation. It becomes critical during ischemic stroke, when large extracellular ATP release from dying and stressed cells increases cytotoxicity and neuroinflammation. CRAC channel–mediated calcium signaling may mediate purinergic responsiveness and the mobilization of processes required for efficient CD39-dependent ATP hydrolysis (Figure 5). In *Stim* cKO mice, impaired calcium influx could therefore compromise both vasoactive adenosine production and extracellular ATP clearance, potentially contributing to the mechanism behind larger infarct volumes and lower ADC values observed 48 hours after stroke in our experiments.

The residual calcium signaling is also relevant to interpreting the effects of our genetic manipulation. Although *Stim1* and *Stim2* mRNA were reduced by approximately two-thirds in isolated microglia, CSD-associated calcium activity decreased by only 33%. Residual responses could reflect incomplete deletion, intracellular calcium release, or STIM-independent influx through channels such as P2X7 and P2X4 (Verma et al., 2017; Ye et al., 2017). We therefore cannot interpret these calcium responses as a direct measure of CRAC conductance, or as proof that microglial calcium signaling was abolished. Nevertheless, the intervention was sufficient to alter infarct evolution. Other work shows that dampening different parts of the microglial calcium machinery can lead to different outcomes. Deleting P2X4, or inhibiting Trpv4, Trpv2, or Kv1.3, has been reported to protect the ischemic brain (Ding et al., 2025; Fomina et al., 2021; Ren et al., 2024; Verma et al., 2017). Yet loss of *Stim1* and *Stim2* worsened injury in our hands. The effect seems to depend on the pathway and the cellular context, not on the overall size of the calcium response.

We used 15-month-old mice, five months after tamoxifen induction, because stroke incidence rises steeply with age (Kelly-Hayes, 2010). Aging and reproductive senescence may change ischemic vulnerability and microglial responses, and earlier work has found sex differences in both. Microglia differ between males and females in density, transcriptome, and inflammatory tone starting from development, and female microglia can restrict ischemic injury even after transplantation into a male brain (Kerr et al., 2019; Liu et al., 2009; Manwani et al., 2013; Ugidos et al., 2022; Villa et al., 2018). The larger 48 h infarcts in cKO mice reached significance in males but not in females. This does not show a sex difference, because the genotype × sex interaction was not significant (2.62-fold, 95% CI 0.78-8.74, p = 0.12) and the female comparison had only 39% power to detect the effect seen in males. Hence, our data cannot distinguish a true sex difference from low statistical power. Also, the 6 h cohort was entirely male, leaving the early perfusion deficit and later growth pattern untested in females.

This study has several limitations arising from endpoints that were not assessed. *Stim1* and *Stim2* mRNA levels in isolated microglia fell by about two-thirds, but protein levels were not quantified. We did not measure cytokines, NFAT activity, microglial polarization, vascular reactivity, or behavior, so a shift in inflammatory balance and its effect on outcome are inferred rather than shown. The model was also permanent distal MCAo read at 48 h, so our data describe acute cortical infarction, not reperfusion injury or later repair.

Taken together, our findings identify STIM-dependent signaling as a protective mechanism during acute cortical ischemic injury. The central result is a sequence of synergistic events. Early strokes at 6 h had comparable lesion volume and edema, but perfusion was already lower in cKO mice. Subsequently, cKO infarcts expanded between 6 and 48 h, with no detectable difference in gross arterial anatomy or the angiographic extent of occlusion. This pattern suggests that cKO mice were less able to preserve vulnerable tissue. Calcium-dependent inflammatory signaling can be injurious (Grinberg et al., 2013; Mizuma et al., 2019). Attenuating this pathway might have been expected to help, yet it was not protective here.

Therefore, reducing microglial STIM-dependent signaling may also impair functions that sustain ischemic tissue. Going forward, the challenge is to reduce damaging inflammation without losing vascular and purinergic support.

## Abbreviations

ADC: apparent diffusion coefficient
ADP: adenosine diphosphate
AMP: adenosine monophosphate
ANOVA: analysis of variance
Arg-1: arginase 1
ASL: arterial spin labeling
ATP: adenosine triphosphate
AUC: area under the curve
Ca²⁺: calcium ion
CBF: cerebral blood flow
cKO: conditional knockout
CRAC: calcium-release activated calcium channels
CSD: cortical spreading depolarizations
CSF1R: colony stimulating factor 1 receptor
DICTs: damage-induced calcium transients
dMCAo: distal middle cerebral artery occlusion
DWI: diffusion-weighted imaging
EPI: echo-planar imaging
ER: endoplasmic reticulum
FAIR: flow-sensitive alternating inversion recovery
GFAP: glial fibrillary acidic protein
IL-1β: interleukin 1 beta
IL-6: interleukin 6
IL-10: interleukin 10
MCAo: middle cerebral artery occlusion
MRA: magnetic resonance angiography
MRI: magnetic resonance imaging
NADH: nicotinamide adenine dinucleotide
NFAT: nuclear factor of activated T cells
NVU: neurovascular unit
PBS: phosphate-buffered saline
PC::G5-tdT: Polr2a-based GCaMP5G and tdTomato Cre-dependent reporter
pMCAo: proximal middle cerebral artery occlusion
PWI: perfusion-weighted imaging
RNA: ribonucleic acid
SD: standard deviation
SOCE: store-operated calcium entry
STIMs: stromal interaction molecules
TGF-β: transforming growth factor beta
TNF-α: tumor necrosis factor alpha
TRP: transient receptor potential
TTC: 2,3,5-triphenyltetrazolium chloride
TurboRARE: rapid acquisition with relaxation enhancement

## Author contributions

K.A.S. and P.T. conceptualized the study and developed the mouse model. L.L., C.V., and K.A.S. performed calcium imaging experiments. M.P. generated calcium signal traces, and M.F.S. analyzed them with assistance from Z.W. C.V. and K.A.S. performed proximal MCAo. P.N. performed distal MCAo surgeries with assistance from K.A.S., N.J., and Z.W. H.L. isolated microglial cells and performed RT-PCR. M.R.H., K.A.S., and C.V. performed MRI acquisition. N.J., Z.W., and A.Z. performed MRI analysis and segmentation under the direction of R.T.K., M.A.J. and G.W.M., who also optimized MRI sequences. P.N., N.J., and K.A.S. performed TTC histology, and Z.W. analyzed the histology data. C.Y.K. provided expertise and resources in stroke and glial biology. K.A.S., N.J., Z.W., P.N., and P.T. designed figures and wrote the manuscript. G.W.M., C.Y.K., and P.T. edited the manuscript. All authors reviewed and approved the final manuscript.

## Acknowledgments

We thank Dr. Stefan Feske for kindly providing the *Stim* conditional mouse strains used in this study. We are grateful to the University of Virginia Advanced Microscopy Facility for support with image data analysis and to the Molecular Imaging Core for help with MRI acquisition.

## Data and code availability

Per-animal imaging measurements, lesion and contralateral region masks, per-slice ADC and per-pixel ASL fits, as well as raw DICOM data and the Python and MATLAB analysis code are available from the corresponding authors on request.

---

**Figure S1.**
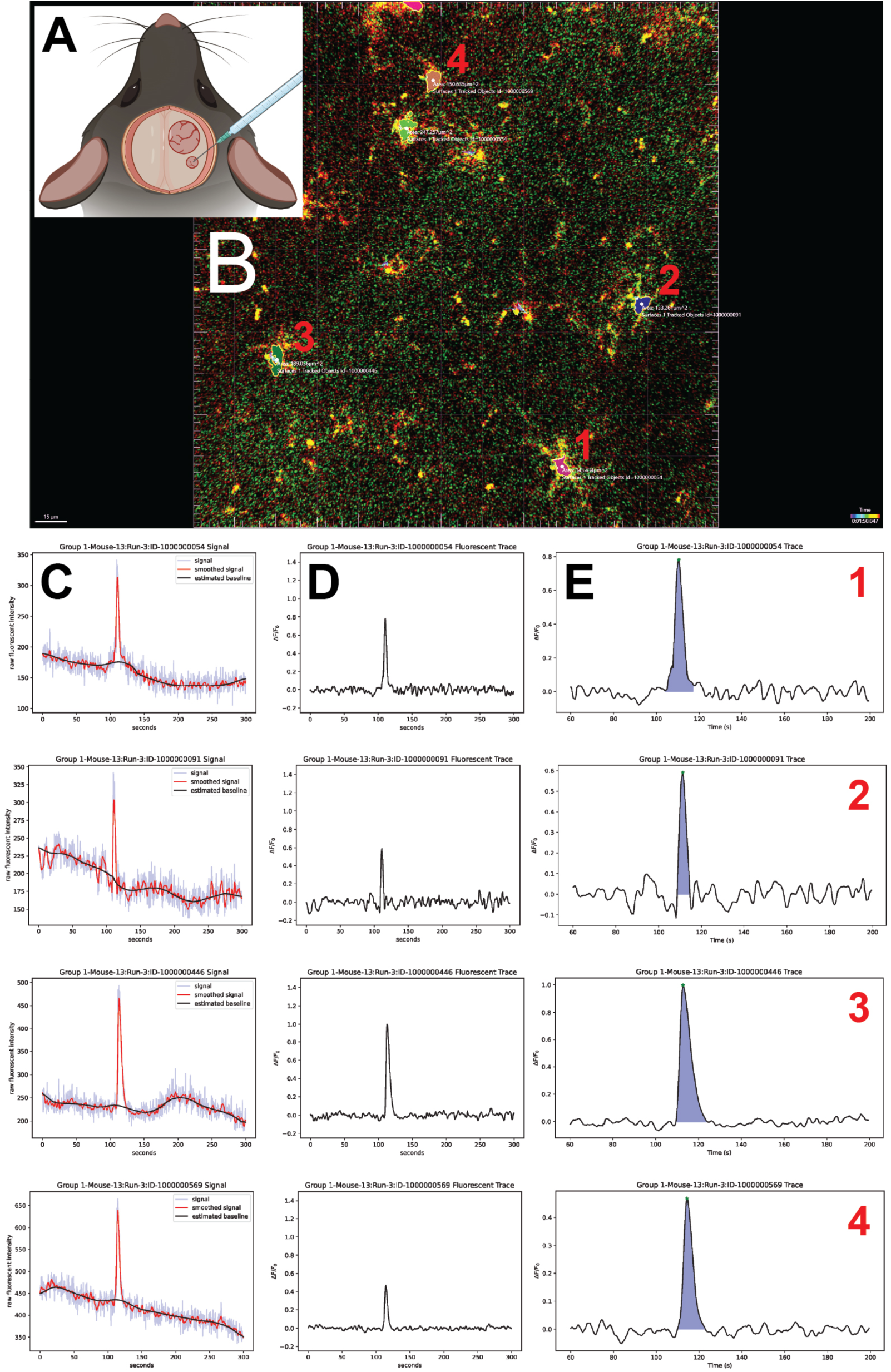
In vivo two-photon imaging and analysis of KCl-evoked microglial calcium transients. **(A)** Schematic of the preparation. A cranial window over the right parietal bone provides optical access to the somatosensory cortex, while cortical spreading depolarization (CSD) is induced through a separate burr hole drilled posteriorly over the visual cortex. CSDs were triggered with 100 nL of 1 M KCl delivered via a glass micropipette. Diagram created with BioRender. (B) Representative merged field of view from a control mouse, showing tdTomato (red) and GCaMP5G (green) fluorescence [Olympus FVMPE-RS, 25x 1.05 NA, 2x zoom, 920 nm, 2.5 Hz]. Microglial somata were segmented and tracked over time in Imaris 9.5 using the tdTomato signal, which reports cell morphology independently of calcium. In this example, four tracked surfaces (1–4) are outlined, each annotated with its cross-sectional area and Imaris tracked-object identifier. Scale bar, 15 µm. (C–E) Signal processing and measurement for the four cells outlined in B. Rows 1–4 correspond to cells 1–4. (C) Mean GCaMP5G intensity within the tracked soma (light blue), the same trace after mean and Wiener filtering (red), and the time-varying baseline F₀(t) estimated by median and Wiener filtering (black). (D) The resulting ΔF/F₀ trace, [F(t) − F₀(t)]/F₀(t), across the full 300-s recording; KCl was injected at t = 60 s. (E) The same trace over the 60–200 s measurement window, showing the three parameters extracted from every transient: peak amplitude (marker), response duration (the interval between the zero crossings bracketing the peak) and area under the curve (shaded). Group data derived from these measurements are presented in Figure 2G–I.

**Figure S2.**
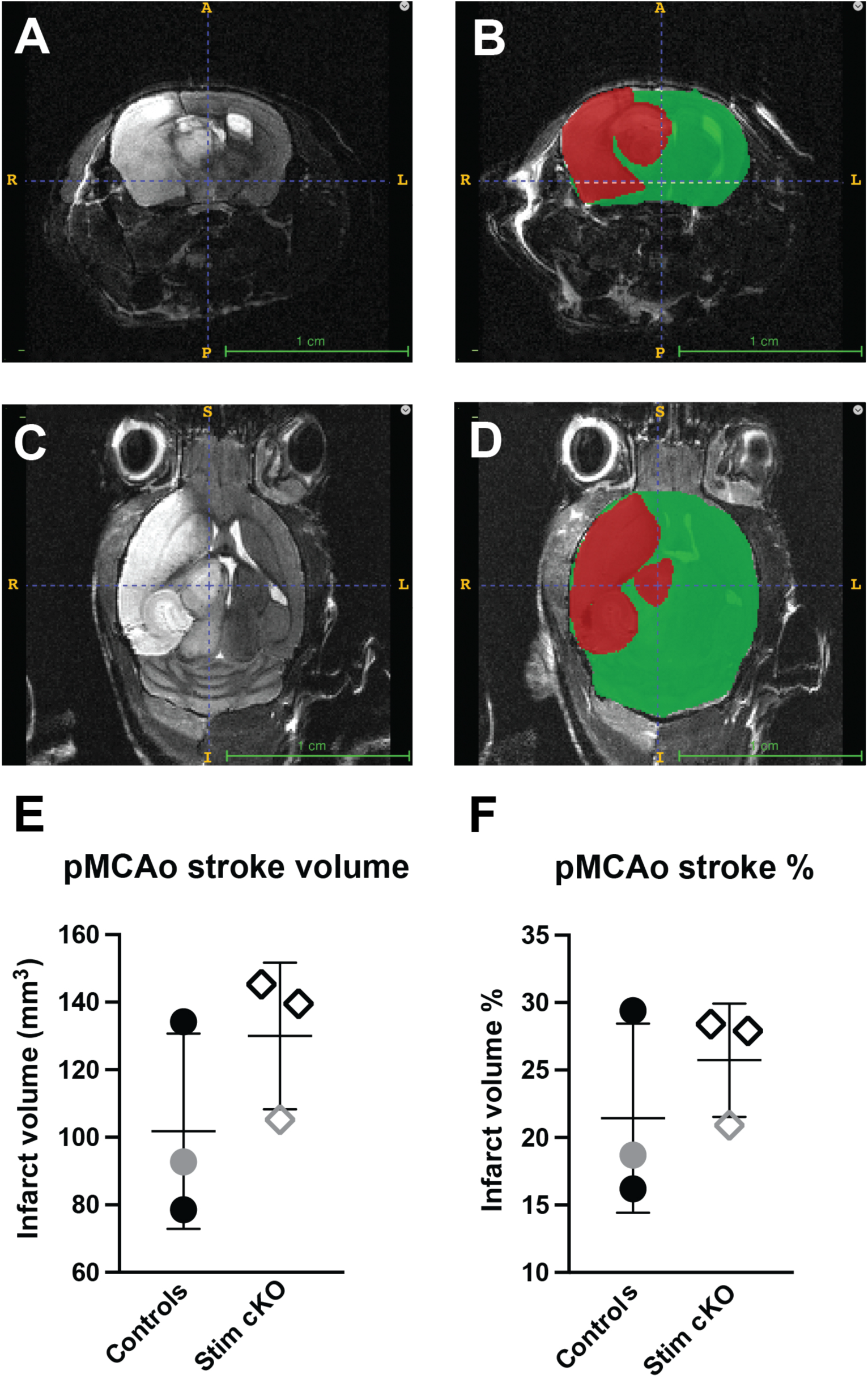
Infarct volume 48 h after proximal middle cerebral artery occlusion in an initial pilot cohort. Ischemic injury was assessed 48 h after pMCAo on a 9.4T scanner using T2-weighted imaging. (A–D) Representative T2-weighted images from a single Stim mutant mouse, shown without overlay (A, C) and with the segmentation masks applied (B, D), in the horizontal (A, B) and coronal (C, D) planes. The infarct (red) and the whole brain (green) were traced manually on every slice using ITK-SNAP (Yushkevich et al., 2006); where the masks overlap, the infarct label is displayed on top. Orientation labels (R, right; L, left; A, anterior; P, posterior; S, superior; I, inferior) and the 1-cm scale bars are those generated by ITK-SNAP. (E, F) Infarct volume (E) and infarct volume as a percentage of total brain volume (F). Volumes were calculated separately from the coronal and horizontal 2D T2 series, and the two measurements were averaged to give the value plotted for each animal. Each group contained two males and one female; male symbols are black and female symbols are grey. Bars indicate mean ± SD. Mean infarct volume was 101.8 ± 28.94 mm³ in controls and 130.0 ± 21.67 mm³ in *Stim* cKO, corresponding to (21.4 ± 7.0)% and (25.7 ± 4.2)% of total brain volume, respectively. n = 3 per group; the differences were not statistically significant.

**Figure S3.**
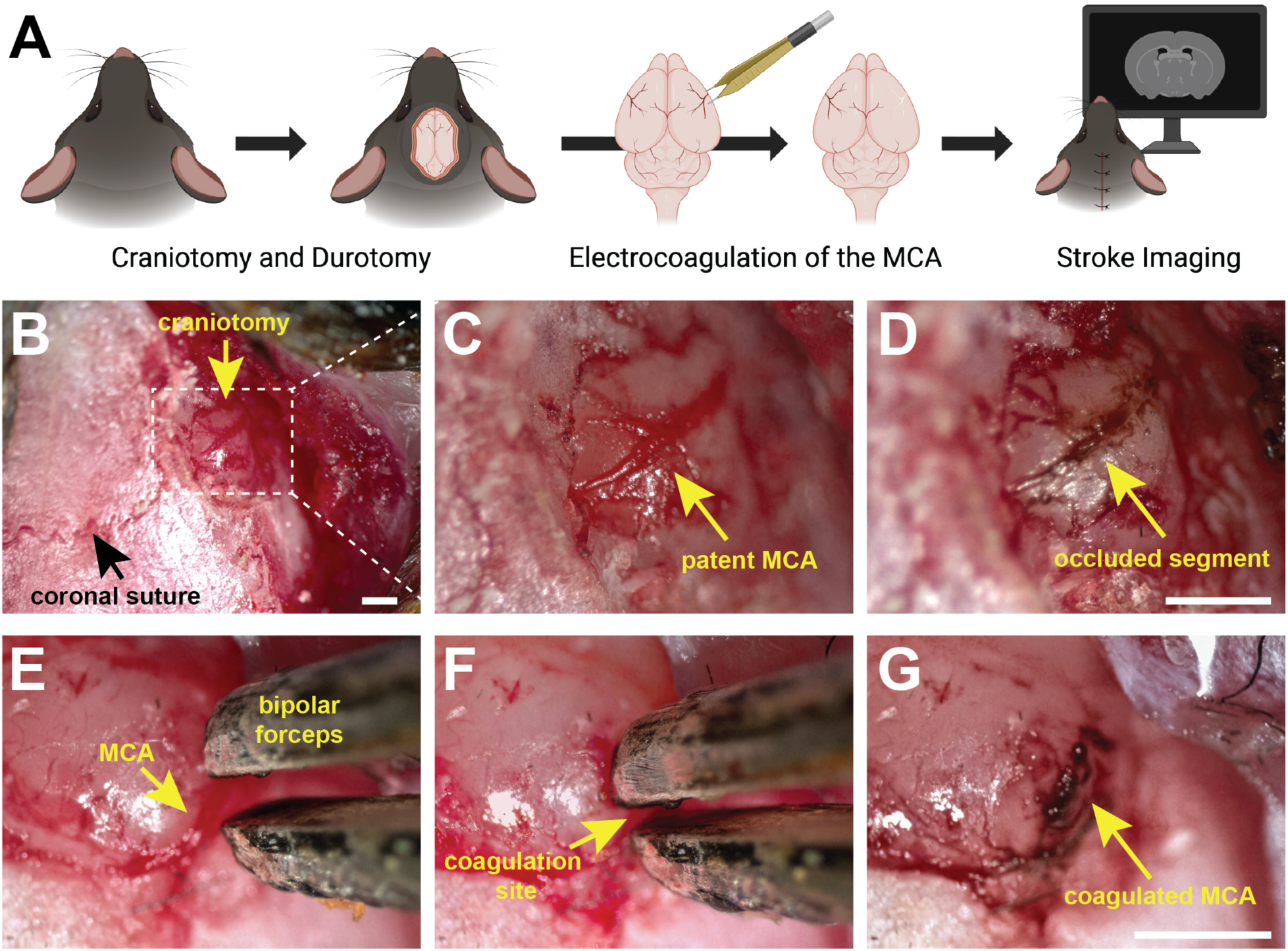
Permanent distal middle cerebral artery occlusion (dMCAo) by bipolar electrocoagulation. (A) Schematic of the procedure. A craniotomy is performed over the temporal bone, and the meninges are opened to expose the distal middle cerebral artery (MCA). The artery is then permanently occluded by bipolar electrocoagulation. The scalp is closed, and infarct development is assessed by MRI. Created with BioRender. (B–D) Intraoperative views of the exposure and occlusion from one representative animal. (B) Lateral skull surface after reflection of the temporalis muscle, showing the craniotomy (yellow arrow) and the coronal suture (black arrowhead). The dashed box marks the field shown in C and D. (C) The distal MCA within the craniotomy before occlusion; the artery is patent and blood-filled (yellow arrow). (D) The same vessel immediately after electrocoagulation. The occluded segment is dark and no longer perfused (yellow arrow). C and D were acquired at the same magnification, cropped and aligned on the medial margin of the craniotomy. (E–G) The occlusion step at higher magnification in a second animal. (E) The bipolar forceps are advanced through the craniotomy toward the exposed MCA (yellow arrow). (F) The forceps are positioned astride the artery at the point of coagulation (yellow arrow). (G) After the procedure, the coagulated arterial segment remains as a dark, retracted stump (yellow arrow), and the vessel is no longer perfused. Scale bars, 1 mm. The bar in D applies to C and D; the bar in G applies to E–G.

**Figure S4.**
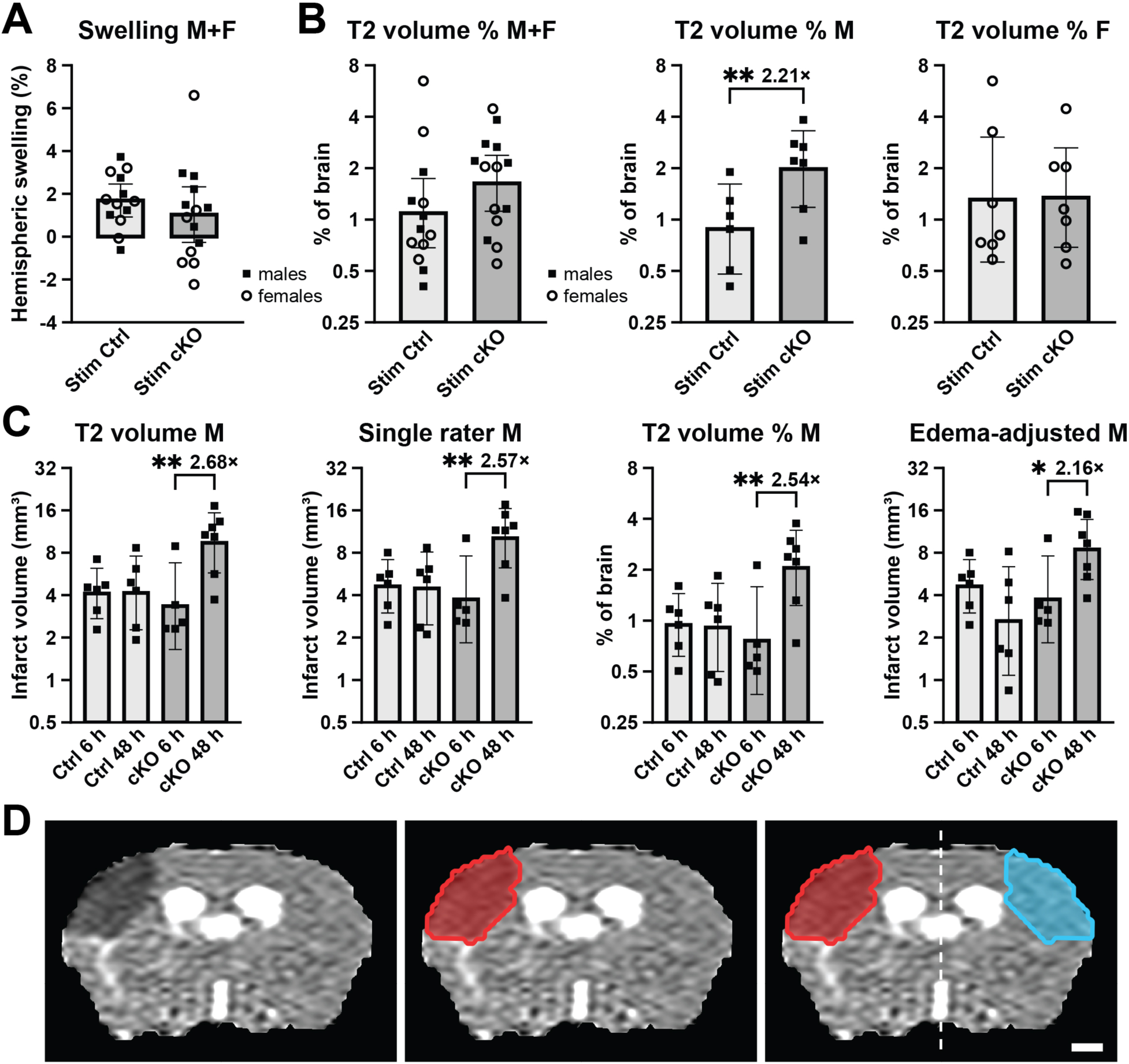
Hemispheric swelling, relative infarct volume, infarct growth from 6 h to 48 h, and the contralateral ADC reference region after dMCAo. (A) Hemispheric swelling. On every coronal T2 slice, the traced brain was divided at the mid-sagittal line. The line was fitted slice by slice by reflection symmetry of the brain outline and T2 intensity. Swelling is the excess volume of the ipsilateral over the contralateral hemisphere, as a percentage of the contralateral volume, within the infarcted slab (infarct slices ± 1 mm). Each value is the mean of the two investigators’ coronal tracings. Values can be negative, so the axis is linear, bars show the arithmetic mean, and error bars show its 95% confidence interval. Across the cohort, the infarcted hemisphere was enlarged by (1.35 ± 1.84)% (20 of 27 mice; p = 0.009, paired t-test of the infarcted slab against lesion-free slices). Swelling did not differ between genotypes (*Stim* Ctrl 1.69%, *Stim* cKO 1.03%; p = 0.35, unpaired Welch t-test). n = 27 (13 control, 14 cKO). (B) Infarct volume as a percentage of total brain volume, for both sexes pooled (left), males (center) and females (right). The animals and tracings are those of Figure 4B–D (each symbol is the mean of four tracings: two investigators × two planes). The y-axis is logarithmic; bars show the geometric mean, and error bars show its 95% confidence interval. Relative infarct volume was larger in male mutants (2.21-fold, p = 0.0046) but not in females (0.86-fold, p = 0.77) or in the pooled cohort (1.28-fold, p = 0.46); fold changes and p values are from a gamma GLM with a log link. n = 27 (males: 6 control, 7 cKO; females: 7 per genotype). (C) Infarct growth from 6 h to 48 h in males. Each plot shows control and *Stim* cKO mice at 6 h (Figure 3) and at 48 h (Figure 4); the 6-h cohort was all male and independent of the 48-h cohort. From left: infarct volume as plotted in Figures 3B and 4C (6 h, mean of two coronal tracings; 48 h, mean of four tracings); the coronal tracing of the single investigator who traced both time points; the same as a percentage of brain volume; and the 48-h volume after subtraction of the swelling-related excess volume (A; 6-h volumes uncorrected, as no swelling was detectable at 6 h). Brackets compare 48 h with 6 h within each genotype (gamma GLM with a log link; fold change = 48-h/6-h ratio of means; unadjusted). Mutant infarcts grew 2.68-fold (p = 0.0032), 2.57-fold (p = 0.0047), 2.54-fold (p = 0.0071) and 2.16-fold (p = 0.029), whereas control infarcts did not change (1.08-, 1.02-, 1.03-and 0.70-fold; all p ≥ 0.33). Growth differed between genotypes in every measure (time × genotype interaction in a joint gamma GLM: 2.48-, 2.52-, 2.48-and 3.08-fold; p = 0.033, 0.028, 0.036 and 0.026). The y-axes are logarithmic; bars show the geometric mean, and error bars show its 95% confidence interval. n = 6 control and 5 cKO at 6 h; 6 control and 7 cKO at 48 h. (D) Construction of the contralateral reference region for ADC, shown on the slice with the largest infarct cross-section in a female control mouse. Left, ADC map (trace ADC over the intact diffusion directions; grey scale 0–1.3 × 10^−3^ mm^2^/s). Center, the T2-defined infarct (red), carried onto the diffusion images by registration. Right, the infarct and its mirror image (cyan) across the slice’s mid-sagittal line (dashed), which was fitted automatically by reflection symmetry. Mean ADC in the two regions, pooled over all usable slices containing the infarct, gives the values in Figure 4H–J. The same mirror construction defines the contralateral regions for CBF and % T2 (Figures 3 and 4). Scale bar, 1 mm.

**Figure S5.**
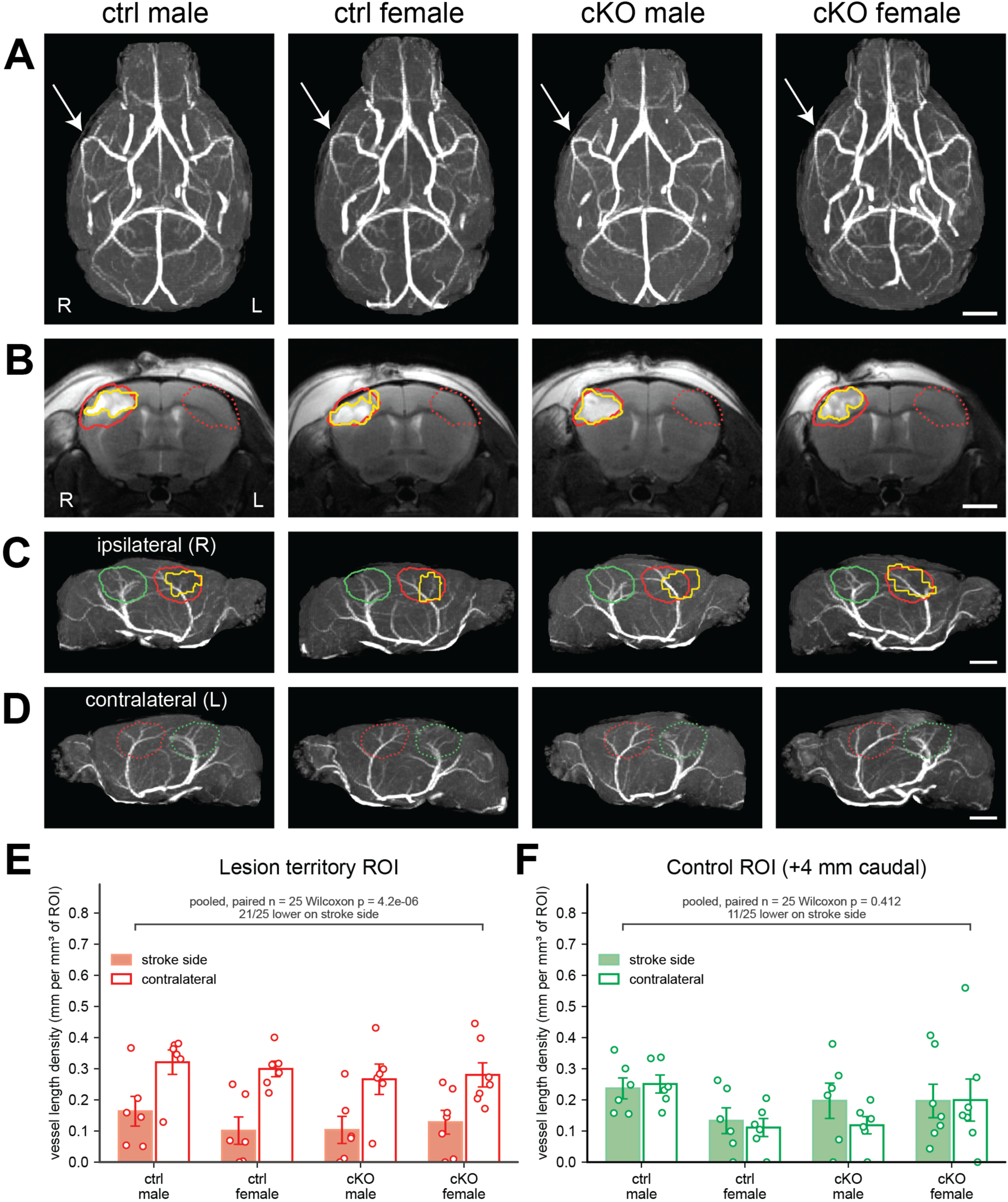
Cerebral angioarchitecture does not differ by sex or *Stim* genotype, and distal MCAo produces a lesion-territory-specific vascular deficit. Time-of-flight MR angiography (TOF-MRA) and T2-RARE, 48 h after distal occlusion of the right middle cerebral artery. Each column is the animal with the median infarct volume in that sex × genotype group. n = 25 (control male 6, control female 6, cKO male 6, cKO female 7). (A) Dorsal maximum-intensity projections of the whole-brain TOF-MRA. Images follow radiological convention, with the right (stroke) hemisphere on the viewer’s left; R and L are marked. Arrows indicate the right middle cerebral artery. The circle of Willis was complete, and both internal carotid arteries were patent in every animal. After registration to a common frame, vessels were segmented and skeletonized, and length, volume, and branch-point count were measured per hemisphere. A ±0.75 mm midline band was left out so that unpaired vessels (basilar artery, azygos anterior cerebral artery, superior sagittal sinus) were not assigned to either side. Right-to-left length ratio was 1.010 ± 0.124 (SD; SEM = 0.025), not different from 1 (p = 0.58). No sex or genotype difference was detected in any hemispheric measure (Holm-adjusted p > 0.05; Kruskal–Wallis on the length ratio across the four groups, p = 0.39). Scale bar, 2 mm. (B) Coronal T2-RARE at the plane of maximal infarct cross-section, matched across columns to within 0.72 mm along the rostro-caudal axis. Yellow, the first rater’s manual edema mask, drawn slice-by-slice on the native coronal T2-RARE and used for all volumetric analyses in this study. Red solid, the fixed lesion-territory ROI. Red dotted, the same ROI mirrored onto the contralateral hemisphere. The fixed ROI is derived from those manual masks and is identical in every animal. Rater’s mask was transformed into the common frame, dilated by 0.4 mm so that the region reaches the pial vessel bed, and the 25 resulting regions were overlaid. Voxels present in at least half the animals (≥ 12 of 25) define the fixed ROI, 12.94 ± 0.54 (SD; SEM = 0.108) mm³ after clipping to each brain. A single fixed region is used because density measured in a lesion-shaped ROI falls as that ROI grows. Scale bar, 2 mm. (C) Ipsilateral (right) sagittal slab projections, 1.25–6.0 mm lateral to the midline, which isolate the middle cerebral artery and its cortical branches. Red, fixed lesion-territory ROI; green, fixed control ROI (the same shape, displaced 4 mm caudally into visual cortex); yellow, this animal’s manual edema mask. Scale bar, 2 mm. (D) Contralateral (left) slab projections, with both ROIs mirrored about each animal’s own midline (dotted). All tissue shown here is uninjured, and the outlines mark the regions against which the ipsilateral measurements are compared. Scale bar, 2 mm. (E, F) Vessel length density, in mm of skeletonized vessel per mm³ of ROI, in the fixed lesion-territory ROI (E) and the fixed control ROI (F). Filled bars, stroke side; open bars, contralateral side. Symbols are individual animals; bars are group mean ± SEM. The bracket gives the pooled within-animal comparison across all 25 animals (two-sided Wilcoxon signed-rank). In the lesion territory, vessel density was 0.124 ± 0.021 on the stroke side versus 0.291 ± 0.019 contralaterally, reduced in 21 of 25 animals (W = 12, p = 4.2 × 10⁻⁶). In the control territory, the two sides did not differ (0.191 ± 0.024 versus 0.171 ± 0.024; 11 of 25; W = 111, p = 0.41). The deficit is therefore confined to the MCA territory and is not a hemispheric bias. Neither panel showed a genotype or sex difference on either side (Mann–Whitney U, all Holm-adjusted p ≥ 0.84). The four sex × genotype groups are shown for display only; the bracketed stroke-versus-contralateral comparison is pooled across all 25 animals, and no per-group paired tests were performed (see Methods).

**Figure S6.**
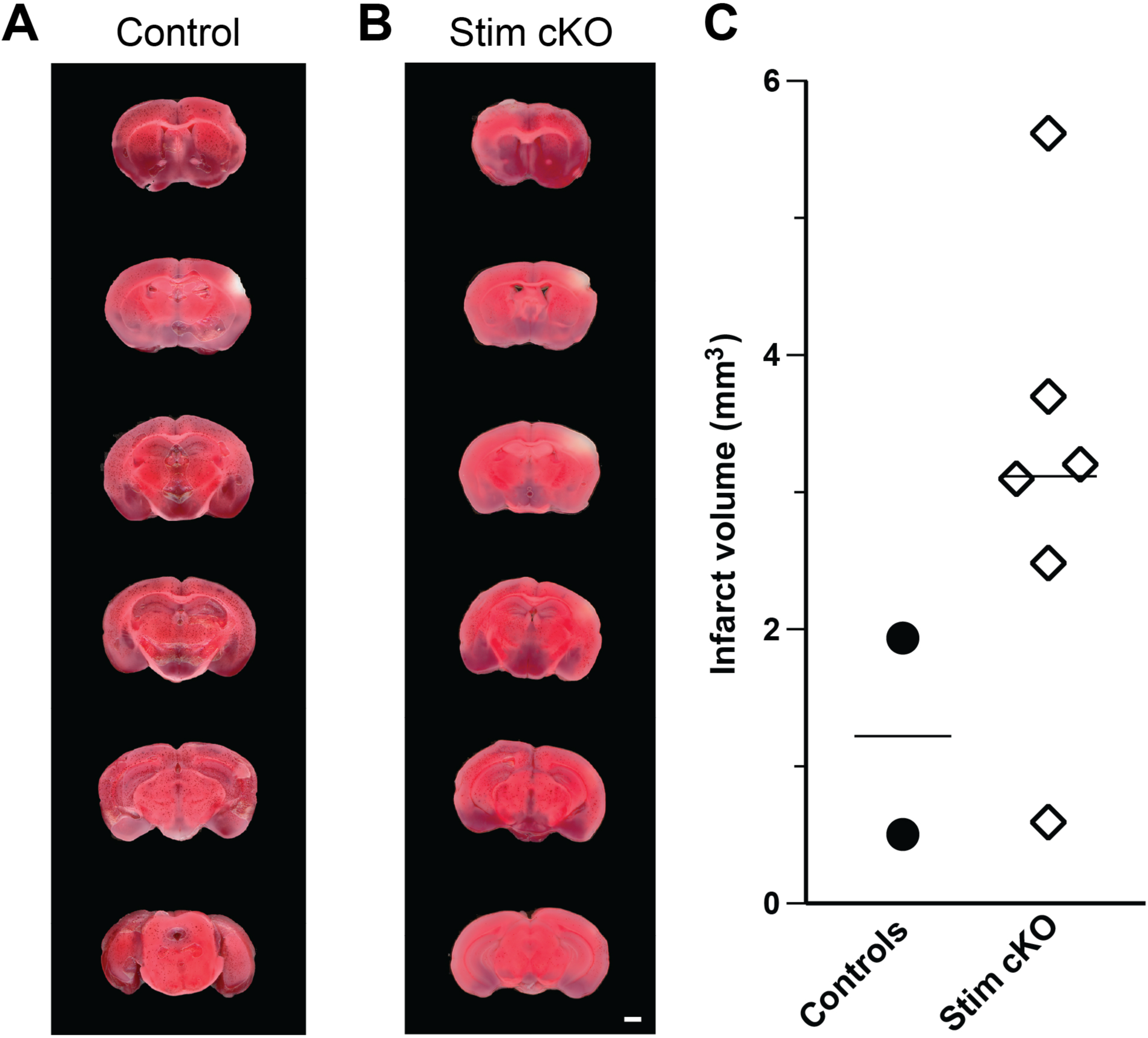
TTC histology is consistent with larger infarcts in male *Stim* cKO 48 h after dMCAo. Immediately after the 48-h MRI session, brains were sectioned into 1-mm coronal slices and stained with 2% TTC, in which viable tissue reduces to red formazan while infarcted tissue remains pale. (A, B) Serial coronal slices from a representative control (A) and a representative *Stim* cKO (B), shown rostral (top) to caudal (bottom) at matched levels; the infarct is the pale region in the right hemisphere, ipsilateral to the occlusion. Scale bar, 1 mm. (C) Infarct volumes summed across slices for individual male mice; horizontal bars indicate group means. Control, n = 2; Stim mutant, n = 6. TTC staining was adequate for quantification in 8 males from 26 brains processed, reflecting incomplete loss of dehydrogenase activity in small dMCAo infarcts. Brains with incomplete or uneven staining were excluded before measurement. Because only two male controls yielded quantifiable sections, no statistical test was performed, and the panel is presented as descriptive support for the MRI data. Mean infarct volume was 1.22 mm³ in controls and 3.12 mm³ in *Stim* cKO, a difference in the same direction as the T2-defined volumes in Figure 4C. Absolute TTC volumes are lower than the corresponding T2 volumes, as expected, because T2 hyperintensity includes vasogenic edema whereas TTC reports loss of dehydrogenase activity.

