## Supplemental Methods for "Microglial Stim1/Stim2 deletion reduces calcium signaling but promotes infarct expansion after ischemic stroke"

**S1. MRI data and acquisition**

The 6-h cohort was imaged in two sessions (2024-03-30 and 2024-05-04) and the 48-h cohort in six (2024-01-08, 2024-01-09, 2024-01-15, 2024-01-16, 2024-01-29, 2024-01-30). Each mouse was scanned once, in a single session containing T2-weighted, diffusion-weighted and ASL acquisitions; time-of-flight angiography was added at 48 h. In total, 57 diffusion scans from 46 mice and 49 FAIR scans from 46 mice were acquired. Acquisition parameters, read from the DICOM headers of every scan, are summarized in Table SM1.

**Table SM1.** MRI acquisition parameters (ranges over all scans). DWI: b = 22–24 s/mm^2^ (reference) and 690–706 s/mm^2^ along three orthogonal directions; field of view 15 mm at 48 h and in the first 6-h session, 20 mm in the second. FAIR: inversion times 12–2300 ms (13 values); 6-h scans had 4000, 6000 and 8000 ms in addition. All scans used a volume transmit coil and a mouse-brain array receive coil (ParaVision 360.3.4).

|  | **T2 (coronal)** | **T2 (axial)** | **DWI** | **FAIR ASL** |
| --- | --- | --- | --- | --- |
| Sequence | 2D TurboRARE | 2D TurboRARE | segmented spin-echo EPI (DtiEpi) | FAIR, segmented EPI |
| TR (ms) | 2500–2746 | 1082–2500 | 2000 | 10022 |
| TE (ms) | 34.4 | 34.4 | 16.9 | 11.1 |
| Refocusing (T2) / excitation angle (°) | 126–180 | 126–180 | 90 | 90 |
| Echo train / segments | 8 | 8 | 15 / 4 (62.5% of 96 lines) | 31 / 2 (77.5% of 80 lines) |
| Bandwidth (Hz/pixel) | – | – | 3472 | 4261 |
| Averages | 4 | 4 | 4 | 1 |
| Matrix; field of view | 220 × 220; 20 mm | 220 × 220; 20 mm | 96 × 96; 15 or 20 mm | 80 × 80; 18 mm |
| Slices × thickness | 30–33 × 0.5 mm | 13–18 × 0.5 mm | 30–33 × 0.5 mm | 1 slab × 1 mm |
| Duration (min) | 4.5–4.9 | 1.9–4.5 | 2.1 | 9.4 (13 TIs) or 12.6 (16 TIs) |

Diffusion scans were repeated in 9 mice (DMCAO 101, 103, 107, 202, 203, 204, 205, 206, 207), including left-right phase-encoded repeats in five 6-h mice acquired to test distortion. FAIR scans were repeated in 3 mice (DMCAO 5, 202, 206). The scan used was chosen before any lesion value was computed (S3.6, S4.6). Vendor (ParaVision) parameter maps were available for 43 diffusion scans and 40 FAIR scans; none were produced in the second 6-h session.

At 48 h, angiography followed DWI, and the first ASL scan started 17.8–22.6 min (median 18.4) after the start of DWI. At 6 h, the first ASL scan after the DWI scan used started a median of 3.9 min later (range 2.7–22.1 min; the longest intervals, in DMCAO 101 and 107 and 103, followed a DWI scan that was then repeated). In two mice (DMCAO 201, 202) an ASL scan was also acquired before DWI. These intervals matter where circulation failed during a session (S4.5).

**S2. Lesion masks and inter-rater agreement**

Infarct and whole brain were traced in ITK-SNAP on every slice. At 6 h, two investigators (N.J., Z.W.) traced the coronal stacks. At 48 h, two investigators (N.J., C.V.) traced both the coronal and the axial stacks. A third investigator (A.Z.) traced most animals independently; these tracings were used only for the agreement audit below. Their files carried no orientation, and their placement on the T2 grid was recovered from the export geometry. Mice whose infarct was at or below the detection limit of the T2 sequence (about 0.5% of brain volume) were excluded before analysis: 7 of 34 at 48 h, and at 6 h DMCAO 105 (0.35%) and 101 (0.19%).

**Table SM2.** Agreement between coronal infarct tracings. ICC(A,1): absolute-agreement intraclass correlation for a single tracing.

| **Cohort** | **Pair** | **Mice** | **Dice, median [IQR]** | **ICC(A,1), log volume** |
| --- | --- | --- | --- | --- |
| 6 h | N.J. – Z.W. | 13 | 0.77 [0.75-0.82] | 0.90 |
| 6 h | N.J. – A.Z. | 12 | 0.44 [0.37-0.62] | 0.53 |
| 6 h | Z.W. – A.Z. | 12 | 0.42 [0.36-0.56] | 0.68 |
| 48 h | N.J. – C.V. | 33 | 0.86 [0.83-0.90] | 0.98 |
| 48 h | N.J. – A.Z. | 26 | 0.71 [0.60-0.84] | 0.76 |
| 48 h | C.V. – A.Z. | 26 | 0.73 [0.60-0.85] | 0.83 |

N.J. drew the largest outlines (volume 1.16× and 1.29× the per-mouse mean of all tracings at 48 h and 6 h; A.Z. 0.77× and 0.74×) and captured nearly the whole consensus lesion estimated by STAPLE (Warfield et al., 2004; sensitivity 0.99 at 48 h and 0.99 at 6 h). N.J. was also the only investigator who traced both cohorts. These coronal tracings therefore defined all lesion regions for ADC, CBF, relative T2 signal and swelling; the second investigator's tracings were used for sensitivity analyses (S7). Infarct volumes in the figures are the means of all tracings by the two investigators (two at 6 h, four at 48 h).

**S3. Diffusion processing**

*S3.1 Images and ADC.*

Each Enhanced MR DICOM file holds the reference block and the three diffusion blocks for every slice, identified by the first dimension index; slices were ordered by position along the slice normal, and stored values were rescaled to physical units. The b-value of each block is the trace of its b-matrix, which includes the contribution of the imaging gradients. ADC was computed as the trace estimate, Σ ln(S_0_/S_i_) / Σ (b_i_ − b_0_), over the intact directions (Basser et al., 1994). The trace estimate was 0.5–3.3% higher than the fit (median 0.9%), equally in lesion and contralateral regions.

*S3.2 Registration to the T2 images.*

The reference image of each DWI scan was registered to the T2 stack on which the masks were drawn, blurred to DWI resolution. A DWI voxel at local position x maps to the T2 world position R(x + t + e_PE_ d_PE_(x)) + o, where R and o are the scanner geometry, t is a 3-D translation (head motion) and d_PE_ is a smooth displacement along the phase-encoding direction e_PE_ (EPI distortion; Jezzard & Balaban, 1995), modeled as a cubic B-spline with knots every 2.5 mm in-plane and 5 mm across slices. The translation was found on a grid (±0.30, ±0.30 and ±0.25 mm) and refined by Nelder–Mead search; the B-spline coefficients were then fitted by damped Gauss–Newton optimization (up to 40 iterations, bending penalty 0.05, ridge 0.002). The cost was the normalized cross-correlation within the brain (dilated by two voxels), and each stage was kept only if it improved the correlation. Head motion between series was 0.10 mm (median; maximum 0.75 mm). The correlation rose from 0.82 with the scanner geometry to 0.88. The 95th percentile of the phase-encoding displacement in the brain was 0.04 mm (median; maximum 0.15 mm), below one pixel. On synthetic data with a known distortion, the model recovered its pattern (r = 0.88) but only 48–76% of its size, so these displacements are lower bounds. The size of the correction did not matter: removing the fitted displacement or doubling it changed the 48-h relative-ADC comparison only from 0.878-fold (p = 0.0078) to 0.885-fold (p = 0.012) and 0.874-fold (p = 0.0057), and individual values by at most 2.2 percentage points.

*S3.3 Partial-volume sampling.*

Each DWI voxel was represented by 4 × 4 × 3 points placed around its registered center, and the label of the T2 mask was read at each point (nearest neighbor). The brain and lesion fractions are the proportions of points labeled brain or lesion. Voxels with a brain fraction ≥ 0.5 formed the brain; brain voxels with a lesion fraction ≥ 0.5 formed the lesion region.

*S3.4 Motion dropouts.*

The diffusion images were acquired in four segments without respiratory gating or navigator correction. Bulk motion during the diffusion gradients gives each segment a different phase error, and the signal of a whole slice can cancel in one direction (Anderson & Gore, 1994). Dropouts were detected in normal-appearing tissue (brain minus the lesion dilated by three voxels; slices with at least 150 such voxels): a direction was flagged when the natural logarithm of its median diffusion-weighted/reference ratio deviated by more than 0.10 from its five-slice running median, and the reference image was flagged in the same way on its own median signal. The threshold corresponds to about the 99th percentile of the scans without dropouts. A slice was used when its reference image and at least two directions were intact. Of the lesion slices in the selected scans, 202 of 204 were usable at 48 h and 60 of 68 at 6 h (Table SM3).

**Table SM3.** Motion dropouts by session (coronal scans). At 48 h, 9 mice had ≥ 10% of direction-slices flagged.

| **Session** | **Scans** | **Direction-slices flagged, median (range)** | **Scans with reference dropouts** |
| --- | --- | --- | --- |
| 48 h | 33 | 2.3% (0.0–23.8%) | 3 |
| 6 h 03-30 | 9 | 38.3% (16.0–49.3%) | 7 |
| 6 h 05-04 | 14 | 1.2% (0.0–4.9%) | 0 |

*S3.5 Regional values.*

In every usable slice, ADC was computed from the intact directions only, and the lesion and contralateral regions (S5) were always evaluated on the same slices and directions. Voxels with ADC ≤ 0 or > 1.5 × 10^−3^ mm^2^/s were excluded as cerebrospinal fluid. Lesion and contralateral ADC are voxel means pooled over all usable slices containing both regions. Each animal required at least two usable lesion slices covering at least 50% of the lesion voxels; the smallest values among the analyzed animals were 2 slices and 65%.

*S3.6 Choice of scan and phase-encoding direction.*

Among repeated scans, the order of preference was coronal orientation, dorso-ventral phase encoding, the most usable lesion slices, the fewest dropouts and the best registration. The five 6-h mice scanned with both phase-encoding directions agreed in normal tissue within 2.2–3.9% per slice and hemisphere, but left-right encoding carried more Nyquist ghosting (5.2% versus 1.6%) and diluted thin cortical lesions, so dorso-ventral encoding was used throughout.

*S3.7 Circulation at the time of DWI.*

Global loss of circulation lowers ADC throughout the brain within minutes. The whole-brain normal-tissue ADC therefore served as a lesion-independent check (48-h median 7.13 × 10^−4^ mm^2^/s). It was 35% low in one mouse (DMCAO 31), which was excluded from the ADC analysis, and normal in the two other 48-h mice without perfusion at the later ASL scan (DMCAO 5 and 12; 6.66 and 6.89).

**S4. Perfusion (FAIR ASL) processing**

Frames are labeled as slice-selective or non-selective inversion, and the inversion time of each frame is stored in its header; stored values were rescaled to physical units. All acquired inversion times were used: 16 at 6 h (12–8000 ms) and 13 at 48 h (12–2300 ms). The three long TIs of the 6-h protocol sample the fully recovered signal, which constrains M0 and T1 in each pixel. Because the 48-h protocol lacked them, absolute CBF values are not strictly comparable between the two time points, and genotypes were compared only within a time point. On the 6-h scans, restricting the fits to the 13 shared TIs raised contralateral CBF by a median of 4% (range 0–6%) and lesion CBF by a median of 16% (range −32% to +93%; Figure 3 mice).

*S4.1 Mode and regional values.*

In each pixel, S(TI) = |M0 [1 − 2 exp(−TI/T1)]| was fitted separately to the slice-selective and the non-selective series. M0 is linear and was eliminated analytically; T1 was found on a logarithmic grid (400 values, 50–8000 ms) and refined by golden-section search (45 steps). CBF = 6000 · λ · (T1,non/T1,blood) · (1/T1,sel − 1/T1,non) ml/100 g/min, with T1 in seconds (Belle et al., 1998; Kober et al., 2008), λ = 0.9 ml/g (Herscovitch & Raichle, 1985; 0.89 ml/g was measured in mice, Leithner et al., 2010) and T1,blood = 2.43 s (2.2–2.4 s reported at 9.4 T; Tsekos et al., 1998; Dobre et al., 2007). Lesion and contralateral CBF are the means of the pixel CBF values in each region (S4.2). Where flow is low, noise makes many pixel values negative (6 h: median 29% of lesion pixels, range 0–48%; 48 h: 21%, range 0–38%). They were retained because discarding or clipping them would bias the mean upward.

*S4.2 Masks on the ASL slab.*

All FAIR scans shared the frame of reference of the T2 stacks, so masks were transferred by scanner geometry. Each FAIR voxel (0.225 × 0.225 × 1 mm) was represented by 5 × 5 × 5 points, and brain and lesion fractions were computed as for DWI (threshold 0.5). The T2 intensity averaged over the slab was used for the slab midline (S5). The slab was tilted by 2.9° (median; maximum 6.9°) from the T2 slices at 48 h.

*S4.3 Cerebrospinal fluid.*

No CSF exclusion was applied to CBF, because the cortical lesions and their mirror images lie away from the ventricles. In the 36 mice analyzed, 1 of 2112 mirror pixels reached CSF-like T2 brightness (more than twice the median T2 signal of brain outside the lesion). A looser threshold (1.5×) flagged 175 pixels in 18 mice; removing them changed mirror CBF by a median of 0.0% (range −14% to +13%).

*S4.4 Absent perfusion.*

Scans in which the contralateral hemisphere had a CBF below 20 ml/100 g/min were excluded (absent perfusion; the next lowest value was 100.4 ml/100 g/min). This removed three 48-h mice (DMCAO 5, 12, 31; contralateral CBF 6.3, 7.9, 2.2), two of which belonged to the Figure 4 set. Their angiograms showed little arterial contrast (1.38, 1.44, 0.94, versus a cohort median of 2.40).

*S4.5 Choice of scan.*

Among repeated FAIR scans, the scan with the best inversion-recovery fit (10th percentile of the brain R^2^) was used, then the highest M_0_ signal-to-noise ratio. In the scans used, the median brain R^2^ was at least 0.993 and the M_0_ signal-to-noise ratio 14.7–201.6.

*S4.6 Quality rules.*

Two rules, defined on the signals and applied to every scan without reference to CBF or genotype, guarded against corrupted images and nearly empty pixels. They were introduced after the first analysis of the 6-h data; Table SM4 gives the lesion comparison without each of them. First, at TIs of 4000 ms and longer, both signals have almost fully recovered, so the region-averaged selective and non-selective signals should nearly coincide (M0 taken from the 13-TI fit of the region's non-selective signal). A long TI was dropped from both series of a scan, for all pixels, when the two differed by more than 5% of M0 in the lesion or contralateral region. In every other 6-h scan they differed by at most 4.7%. One image pair failed: DMCAO 103 at 6000 ms, where the lesion signals differed by −7.8% of M0. In that non-selective image, the signal was raised over a dorsal band that included the lesion. In 27 of 70 lesion pixels the difference exceeded three times the noise, against none at 4000 or 8000 ms and none of 504 lesion pixels in the other ten Figure 3 mice at 6000 ms. Because lesion T1,sel and T1,non differ by less than 1%, this single image lowered the lesion CBF of DMCAO 103 from 16.2 to −3.5 ml/100 g/min. Second, pixels with little signal at the tissue edge or in signal voids give unstable T1 pairs. Pixels whose M0 (non-selective fit) was below ten times the background noise σ were therefore excluded from all regions, with σ estimated from the air outside the head over all images (Rayleigh distribution, σ2 = ⟨m2⟩/2). This removed two lesion pixels in DMCAO 104 and three in DMCAO 205 at 6 h, and two and one contralateral pixels in DMCAO 14 and 38 at 48 h.

**S5. Mid-sagittal line and contralateral region**

The same construction was used for ADC (every DWI slice with lesion and at least 100 brain voxels), CBF (the ASL slab) and relative T2 signal (every coronal T2 slice with lesion). The mid-sagittal line is the line about which the slice is most symmetric. Its score is the soft Dice overlap of the brain fraction with its reflection plus the normalized cross-correlation of the T2 intensity with its reflection; the lesion of any investigator, dilated by two pixels, and its mirror image are left out of the intensity term. Angles within ±15° and offsets within ±6 pixels of the brain centroid were searched (1° and 0.2-pixel steps), then refined (0.1° and 0.05 pixels). The lesion side is the sign of the lesion-weighted mean distance from the line. The contralateral region is the lesion fraction reflected across the line (bilinear interpolation, ≥ 0.5), restricted to contralateral brain and stripped of every voxel that any investigator traced as lesion (Figure S4D).

**S6. Hemispheric swelling**

Swelling was measured on the coronal T2 tracings, in which the brain label includes the ventricles. In each slice, the brain was split at a mid-sagittal line fitted as in S5, except that the lesion, dilated by about 1 mm, and its mirror image were hidden from both the outline and the intensity terms, so that the swollen side could not shift the line. The search ran on images at half resolution (0.18 mm) and the line was applied at full resolution, with pixels crossed by the line split proportionally. Swelling is 100 × (V_ipsi_ − V_contra_)/V_contra_ within the infarcted slab (lesion slices ± 1 mm; slices with < 20 mm^2^ of brain excluded). Lesion-free slices at least 2 mm from the lesion with at least 30 mm^2^ of brain served as reference. Because all infarcts were right-sided, the mean reference asymmetry of each tracing set was subtracted as a constant (48 h: +0.70% for N.J. and +0.78% for C.V.; 6 h: +0.63%). Subtracting each animal's own reference asymmetry instead added noise, because slab and reference asymmetries were uncorrelated. The two investigators' 48-h values agreed (r = 0.81 for swelling; r = 0.87 for the excess volume; mean difference 0.04%). Axial tracings could not be used (r = 0.23 and 0.31 with the coronal values), because head roll is through-plane in horizontal slices and the dorsal convexity changes cross-section fastest with height. Animals whose per-slice reference asymmetry varied by more than 5% (SD) were flagged for visual review but not excluded (5 at 48 h, 4 at 6 h); in the animals inspected (DMCAO 14 and 34), the variation came from reference slices in the cerebellum or olfactory bulbs, not from the infarcted slab. The swelling-related excess volume (baseline-corrected ipsilateral minus contralateral slab volume) was subtracted from each investigator's full 48-h coronal infarct volume for the edema-adjusted comparison in Figure S4C.

**S7. Statistical models and robustness**

Positive, right-skewed quantities (infarct volume, CBF) were compared with gamma generalized linear models with a log link (statsmodels; dispersion estimated from the Pearson χ2; Wald tests), which estimate the ratio of group means directly. CBF values below 1 ml/100 g/min (or 1% for relative CBF) were to be set to 1 before fitting, because the gamma model requires positive values; no value in Figures 3 or 4 required it. ADC measures and relative T2 signal were compared on the raw scale with Welch t-tests; the ratio of means is given with Fieller's 95% confidence interval. Each panel of Figures 3 and 4 contains a single comparison. In Figure S4C, the two within-genotype comparisons are not adjusted, because the pre-specified test of differential growth is the time × genotype interaction. Bars show geometric means with 95% confidence intervals except for swelling (arithmetic mean, because values can be negative).

**Table SM4.** Sensitivity of the 6-h lesion CBF comparison (Figure 3C). One control lesion (DMCAO 207) retained high flow (109.9 ml/100 g/min); all five cKO lesions were below 22 ml/100 g/min. The direction was the same in every analysis, but significance depended on the test, on DMCAO 207, on the masks and on the inversion times.

| **Analysis of 6-h lesion CBF (n = 6 control, 5 cKO)** | **cKO/control (95% CI)** | **p** |
| --- | --- | --- |
| Gamma GLM, Wald z (as in Figure 3C) | 0.31 (0.11–0.89) | 0.029 |
| Gamma GLM, t reference (9 df) | 0.31 | 0.057 |
| Gamma GLM, deviance F test | 0.31 | 0.065 |
| Welch t, raw values | 0.31 | 0.17 |
| Welch t, log values | 0.42 | 0.13 |
| Mann–Whitney U | – | 0.13 |
| Exact permutation, ratio of means (462 splits) | 0.31 | 0.12 |
| Gamma GLM without control DMCAO 207 | 0.52 | 0.12 |
| Leave one mouse out (11 fits) | 0.24–0.52 | 0.0078–0.12 |
| Gamma GLM, second investigator, masks drawn on ASL maps | 0.37 (0.14–0.97) | 0.044 |
| Gamma GLM, second investigator, T2 masks | 0.33 (0.10–1.07) | 0.065 |
| Joint shared-M₀ fit | 0.30 (0.11–0.88) | 0.027 |
| 13 inversion times (those shared with 48 h) | 0.38 (0.13–1.06) | 0.064 |
| Long-TI rule at 4% instead of 5% | 0.31 (0.11–0.89) | 0.030 |
| Long-TI rule off (8%; DMCAO 103 set to 1) | 0.23 (0.07–0.75) | 0.015 |
| No low-signal guard | 0.34 (0.12–0.98) | 0.047 |
| Fit of region-averaged signals, 13 TIs (earlier method) | 0.27 (0.09–0.78) | 0.015 |

**Table SM5.** Infarct growth from 6 to 48 h (Figure S4C). n = 6 control and 5 cKO at 6 h; 6 control and 7 cKO at 48 h (independent cohorts). Within-genotype fold changes from two-group gamma GLMs; interaction from the joint model. For the primary measure, a log-scale linear model gave an interaction of 2.79-fold (p = 0.026), a permutation of time labels within genotype p = 0.043, and leaving out one animal at a time p = 0.0003–0.074.

| **Measure (males)** | **Control, 48 h/6 h (p)** | **cKO, 48 h/6 h (p)** | **Time × genotype (p)** |
| --- | --- | --- | --- |
| Infarct volume (Figures 3B, 4C) | 1.08 (0.78) | 2.68 (0.0032) | 2.48 (0.033) |
| Single investigator, coronal | 1.02 (0.94) | 2.57 (0.0047) | 2.52 (0.028) |
| % of brain volume | 1.03 (0.92) | 2.54 (0.0071) | 2.48 (0.036) |
| 48 h adjusted for swelling | 0.70 (0.33) | 2.16 (0.029) | 3.08 (0.026) |

**Table SM6.** Robustness of the 48-h relative ADC comparison (Figure 4J). Second investigator: C.V.; AND/OR: intersection/union of the two investigators' lesions; ParaVision-type fit: least-squares fit instead of the trace estimate. Welch t-test throughout. Only removing the CSF exclusion weakened the result.

| **Analysis variant (48-h relative ADC, n = 13 + 13)** | **cKO/control (95% CI)** | **p** |
| --- | --- | --- |
| primary (N.J.'s masks) | 0.88 (0.81–0.96) | 0.0078 |
| second investigator's masks | 0.89 (0.81–0.97) | 0.015 |
| N.J. AND second investigator | 0.88 (0.81–0.97) | 0.012 |
| N.J. OR second investigator | 0.88 (0.81–0.96) | 0.0091 |
| dropout-free slices only | 0.88 (0.81–0.97) | 0.015 |
| no dropout handling | 0.89 (0.81–0.97) | 0.014 |
| ParaVision-type fit | 0.89 (0.81–0.97) | 0.014 |
| CSF threshold 1.2e-3 | 0.88 (0.81–0.96) | 0.0066 |
| CSF threshold 2.0e-3 | 0.88 (0.81–0.97) | 0.015 |
| no CSF exclusion | 0.89 (0.79–1.01) | 0.063 |
| voxel median | 0.88 (0.81–0.96) | 0.0075 |
| Excluding DMCAO 5 (n = 25) | 0.88 (0.80–0.96) | 0.011 |
| Including DMCAO 31 (n = 27) | 0.86 (0.79–0.95) | 0.0045 |

**S8. Reproducibility**

For independent recalculation, a workbook lists the per-mouse T2 volumes, ADC and CBF values, the scans used and the exclusions, together with the exact lesion and contralateral regions as NIfTI files on the DWI and ASL grids, the per-slice ADC values with their dropout flags, and the per-pixel ASL fits, the inversion times used and the quality-rule decisions of every scan. Code and these files are available from the corresponding authors on request.
